# Integrating Conformational Dynamics and Perturbation-Based Network Modeling for Mutational Profiling of Binding and Allostery in the SARS-CoV-2 Spike Variant Complexes with Antibodies: Balancing Local and Global Determinants of Mutational Escape Mechanisms

**DOI:** 10.1101/2022.06.17.496646

**Authors:** Gennady Verkhivker, Steve Agajanian, Ryan Kassab, Keerthi Krishnan

**Author notes:** Correspondence; (G.V).

## Abstract

In this study, we combined all-atom MD simulations, the ensemble-based mutational scanning of protein stability and binding, and perturbation-based network profiling of allosteric interactions in the SARS-Cov-2 Spike complexes with a panel of cross-reactive and ultra-potent single antibodies (B1-182.1 and A23-58.1) as well as antibody combinations (A19-61.1/B1-182.1 and A19-46.1/B1-182.1). Using this approach, we quantify local and global effects of mutations in the complexes, identify structural stability centers, characterize binding energy hotspots and predict the allosteric control points of long-range interactions and communications. Conformational dynamics and distance fluctuation analysis revealed the antibody-specific structural stability signatures of the spike complexes that can dictate the pattern of mutational escape. By employing an integrated analysis of conformational dynamics and binding energetics, we found that the potent antibodies that efficiently neutralize Omicron spike variant can form the dominant binding energy hotpots with the conserved stability centers in which mutations may be restricted by the requirements of the folding stability and binding to the host receptor. The results show that protein stability and binding energetics of the SARS-CoV-2 spike complexes with the panel of cross-reactive ultrapotent antibodies are tolerant to the constellation of Omicron mutations. A network-based perturbation approach for mutational profiling of allosteric residues potentials revealed how antibody binding can modulate allosteric interactions and identified allosteric control points that can form vulnerable sites for mutational escape. This study suggested a mechanism in which the pattern of specific escape mutants for ultrapotent antibodies may not be solely determined by the binding interaction changes but are driven by a complex balance and tradeoffs between different local and global factors including the impact of mutations on structural stability, binding strength, long-range interactions and fidelity of allosteric signaling.

## 1. Introduction

The rapidly growing body of structural, biochemical and functional studies established that the mechanism of SARS-CoV-2 infection may involve conformational transitions between distinct functional forms of the SARS-CoV-2 viral spike (S) glycoprotein [1–9]. The S protein consists of conformationally adaptive amino (N)-terminal S1 subunit and structurally rigid carboxyl (C)-terminal S2 subunit, where S1 includes an N-terminal domain (NTD), the receptor-binding domain (RBD), and two structurally conserved subdomains SD1 and SD2 that coordinate protein response to binding partners and regulate the interactions with the host cell receptor ACE2. Conformational plasticity of the SARS-CoV-2 S protein are exemplified by spontaneous transitions from the closed state to the open state accompanied by large scale movements of the RBDs that can spontaneously fluctuate between “RBD-down” and “RBD-up” positions, where binding to the host cell receptor ACE2 preferentially stabilizes the receptor-accessible “up” conformation [1–12]. The cryo-EM experimental tools have been deployed at unprecedented speed to characterize dynamic structural changes in the SARS-CoV-2 S protein, revealing a spectrum and atomic details of the prefusion S conformations that included various forms of the closed “RBD-down” state, “partially open” trimers with only one or two RBDs in the “up” conformation and “open” trimers with all three RBDs in the “up” position [5–12]. Structural changes that accompany SARS-CoV-2 S binding with the ACE2 host receptor demonstrated a well-orchestrated cascade of conformational transitions from a compact closed form weakened after furin cleavage to the partially open states and to the ACE2-bound open form [13,14]. The cryo-EM and tomography tools examined conformational flexibility and distribution of the S trimers in situ on the virion surface showing that the underlying physical mechanism of spontaneous conformational changes between different functional open and closed S states and the intrinsic properties of the conformational landscapes for SARS-CoV-2 S trimers are preserved in different biological environments [15]. Single-molecule Fluorescence (Förster) Resonance Energy Transfer (smFRET) has been an informative biophysical imaging tool for revealing the intrinsically dynamic, heterogeneous nature of the SARS-CoV-2 S trimer on virus particles and capturing dynamics of conformational transitions from the closed state to the receptor-bound open state [16]. These studies suggested that conformational selection and receptor-induced structural adaptation of the S states can work synchronously, providing diverse mechanisms for antibody-induced neutralization by either directly competing with the ACE2 receptor binding or by modulating conformational landscape of the S protein and allosterically stabilizing the S protein in the specific conformation [16]. Biophysical studies demonstrated that conformational dynamics and allosteric regulation of the S protein are intrinsically related, in which allosteric modulation of the RBD equilibrium can be a critical component of the SARS-Cov-2 adaption strategies [17]. Using smFRET this study discerned the temporal prevalence of distinct S conformations in the context of virus particles, presenting the experimental evidence of decelerated transition dynamics from the open state and the increased stability of the S open conformations that facilitate access to the host receptor [17]. Allosteric modulation of the conformational dynamics and ligand-induced population shifts in the SARS-CoV-2 S was quantified using an smFRET imaging assay showing that antibodies that target diverse epitope may allosterically promote shift of the RBD equilibrium toward the up conformation, enhancing ACE2 binding[18]. The variants of concern (VOC’s) with the enhanced transmissibility and infectivity profile include D614G variant [19–21], B.1.1.7 (alpha) [22,23] B.1.351 (beta) [24,25], B.1.1.28/P.1 (gamma) [26], B.1.1.427/B.1.429 (epsilon) variant [27,28], and the recent Omicron variants (B.1.1.529, BA.1, BA.1.1, BA.2, BA.3, BA.4 and BA.5 lineages) [29–31]. Structural studies characterized a significant heterogeneity and plasticity of the SARS-CoV-2 S proteins including B.1.1.7 (alpha), B.1.351 (beta), P1 (gamma) and B.1.1.427/B.1.429 (epsilon) variants showing that the intrinsic conformational flexibility of the S protein variants can be controlled by mutations and determine their ability to evade host immunity and incur resistance to antibodies [32–35].

To understand the antigenic anatomy of the SARS-CoV-2 spike protein, and the molecular mechanisms of SARS-CoV-2 neutralization mediated by the rapidly growing number of potent antibodies, several studies examined the diversity of the binding epitopes in the structures of the antibody complexes and presented a detailed classification of these antibodies into distinct categories [36,37]. The structural principles of these classifications and mechanistic aspects of antibody-mediated neutralization were extensively discussed and summarized in several reviews [38–41]. Combining biophysical and structural data, RBD-directed antibodies can be classified into 7 discrete communities often referred to as RBD-1 to RBD-3, RBD-4 to RBD-5, and RBD-6 to RBD-7 [38]. In this classification, class I neutralizing antibodies are characterized by direct ACE2 competition via binding RBD-up conformations, class II neutralizing antibodies bind RBD-up and RBD-up states, class III antibodies bind outside the immediate RBD-ACE2 interface in both RBD-up/RBD-down forms, and class IV antibodies bind to a cryptic epitope in the RBD-up conformations. Although class I and class II antibodies have significant overlap on their structural footprints, escape mutants at K417, D420, N460, and S475 sites were observed only for class I antibodies. At the same time, class 2 antibodies were escaped by L452R, E484, E490, and Q493 escape mutants that are characteristic for this class. Importantly, these studies established that the RBD-2 class of neutralizing antibodies exhibit the highest potency but suffer from the greatest sensitivity to viral escape, while antibodies targeting RBD-1 and RBD-3 epitopes are characterized by the reduced potency but displaying the greater breadth and tolerance to RBD mutations [38–42]. Combinations and synergistic cocktails of different antibody classes simultaneously targeting the conserved and more variable SARS-CoV-2 RBD epitopes can provide more efficient cross-neutralization effects and yield resilience against mutational escape [43,44]. Some ultra-potent antibodies may function by allosterically interfering with the host receptor binding and causing conformational changes in the S protein that can obstruct other epitopes and block virus infection without directly interfering with ACE2 recognition. Functional mapping of mutations in the S-RBD regions using deep mutational scanning showed that sites of antibody mutational escape are constrained by requirement of the protein stability and ACE2 binding, rationalizing why escape-resistant antibody cocktails often curtail virus evolvability to acquire new RBD sites of immune escape [45–47].

The constellation of Omicron mutations facilitates the evasion of immune responses induced by vaccination and confer resistance to most of the neutralizing antibodies. Recent structural biology studies of the Omicron variant in various functional states and complexes with antibodies suggested that the Omicron variant may have evolved to mediate diverse neutralization escape via multiple mutations and different mechanisms, while enhancing binding affinity with ACE2 using mutational changes in several key energy hotspots [48–53]. Structural and biochemical characterizations of the S Omicron trimer binding with ACE2 and JMB2002 antibody showed that the Omicron spike protein can strengthen binding to the ACE2 receptor, while two JMB2002 antibody molecules able to bind to one RBD-up and one RBD-down protomers effectively neutralizing Omicron [52]. Structural analysis of the SARS-CoV-2 Omicron S protein states showed that unlike other VOC’s implicated in the enhanced viral transmissibility through mutation-induced stabilization of the open S states, the Omicron variant can lead to the increased thermodynamic stabilization of the RBD-down closed state and promote immune evasion by occluding immunogenic sites [53]. The structures of the S Omicron bound to a patient-derived Fab antibody fragment (510A5) showed that antibody binding can be greatly reduced by the Omicron mutations and yet retain strong ACE2 interactions [54]. Using an arsenal of biophysical tools including antigen-based flow-sorting and live virus neutralization assays, a recent study identified four antibodies (A19-46.1, A19-61.1, A23-58.1, B1-182.1) which target the RBD and neutralize SARS-CoV-2 WA-1 original virus strain, while also maintaining potent neutralizing activity against 23 variants, including B.1.1.7, B.1.351, P.1, B.1.429, B.1.526, B.1.529.1, B.1.617.1 and B.1.617.2 VOCs. [55]. The cryo-EM reconstructions for structures of the A23-58.1 and B1-182.1 bound to S-WA1 revealed that the antibodies bind to the S protein with all RBDs in the up position. An impressive structure-functional tour-de-force investigation of the impact of the SARS-CoV-2 S mutations on the binding and neutralization of monoclonal antibodies confirmed that many potent antibodies targeting the spike RBD experienced a significant loss of binding for B.1.1.529 variant [56]. Using functional assays and cryo-EM structures of the S Omicron complexes with a large panel of antibodies, this study identified that only A23-58.1, B1-182.1, COV2-2196, S2E12, A19-46.1, S309, and LY-CoV1404 maintain substantial neutralization against the Omicron variant. Class I B1-182.1 and class II A19-46.1 antibody revealed potent neutralization of B.1.1.529, while class III and IV antibodies that bind outside of the ACE2-binding surface (A19-61.1, COV2-2130, S309, and LY-CoV1404) can tolerate individual B.1.1.529 substitutions [56]. The structure of the ternary complex of the Omicron complex with a combination of B1-182.1 and A19-46.1 antibodies targeting different binding epitopes and trapping 3 RBD-up S conformation suggested a mechanism of the observed synergistic increase in neutralization potency compared with that of the individual antibodies [56]. A recent study identified a neutralizing human monoclonal antibody 87G7 with a broad-spectrum neutralization efficacy achieved via ACE2 binding inhibition mechanism and showing robust neutralizing activity against multiple VOC’s including the Omicron variants [57]. The cryo-EM structure revealed that 87G7 binds the ACE2 receptor binding site by targeting a patch of hydrophobic residues Y421, L455, F456, F486 and Y489 that are highly conserved among SARS-CoV-2 variants and where mutations occur at extremely low frequency (< 0.05%) of human-derived SARS-CoV-2 sequences. A considerable number of residues that are mutated in SARS-CoV-2 variants, including K417N, L452R, S477N, T478K, E484A, E484K, F490S, Q493R are close to the 87G7 core epitope but are unable to elicit mutational escape [57]. Biochemical and functional studies isolated 323 human monoclonal antibodies and showed that a subset of 24 out of 163 RBD-recognizing antibodies potently neutralized all SARS-CoV-2 variants of concern, including Omicron[58]. The cryo-EM structures of a prefusion-stabilized S Omicron trimer and binding assays revealed two highly potent antibodies XGv347 and XGv289 neutralizing B.1.1.7, B.1.351, P.1, B.1.617.2, and B.1.1.539 due to the extensive hydrophobic interactions with F456, Y473, F486 and Y489, and revealing G446S as a single critical mutation site that can confer resistance [58]. Subsequent studies isolated ~ 400 monoclonal antibodies from participants received 3 doses of inactivated vaccine, with a subset of these antibodies exhibiting broad and potent neutralization activities against all SARS-CoV-2 variants [59]. Structural analysis of the most potent XGv051, XGv264, and XGv286 antibodies against Omicron sublineages showed that binding is driven by a network of hydrophobic interactions with the RBD residues Y449, Y453, L455, A475, A484, F486 and Y489 which is characteristic of A23-58.1 and other broadly cross-reactive and ultrapotent antibodies targeting stability-essential RBD sites while being tolerant to the Omicron substitutions [59]. Structure-functional studies also characterized highly potent antibodies G32R7, C98C7, and G32Q4 from each of the RBD classes (RBD-1, RBD-2, RBD-3) that neutralized all five VOCs from Alpha to Omicron BA.1 unveiling the molecular determinants of broad neutralization for class II antibodies [59]. Analysis of the neutralization profiles for a broad panel of antibodies against the Omicron sub-lineages, showed that while most antibodies lost neutralizing activity, some display a unique Omicron escape potential reflecting antigenic difference [60]. The effect of individual and combined mutations that convergently appeared in different lineages was examined showing that the RBD sites more dispensable for binding to ACE2 (particularly E484 and S494) can be hotspots for immune evasion via E484K and S494P modifications across multiple antibodies, and in the combination with mutations that promote ACE2 binding, such as N501Y, can increase escape neutralizing-antibody responses [61]. Besides class I-III antibodies with broad activity against multiple VOC’s, the recent investigation identified two highly conserved cryptic regions on the S-RBD Omicron that are simultaneously and synergistically recognized by bispecific single-domain antibody [62]. The recent pioneering discoveries of broadly neutralizing antibodies that are effective against multiple VOC’s showed that they target highly conserved binding epitopes or conserved cryptic regions, where the antibody-interacting RBD residues are rarely mutated in GISAID database [63,64]. Collectively, the growing number of structural, functional and biophysical studies revealed the diversity of mechanistic scenarios underlying antibody binding and catalogued the RBD escape mutations for a wide range of antibodies revealing distinct signatures of antibody-resistant mutational hotspots. Computer simulations provided important atomistic and mechanistic insights into understanding of the dynamics and function of SARS-CoV-2 glycoproteins. Molecular dynamics (MD) simulations of the SARS-CoV-2 S proteins and mutants detailed conformational changes and diversity of ensembles, demonstrating enhanced functional and structural plasticity of the S proteins [65–77]. All-atom MD simulations of the S-protein in solution and targeted MD simulations of conformational changes between open and closed forms revealed the key electrostatic interdomain interactions mediating protein stability and kinetics of functional spike states [76]. Generalized replica exchange MD simulations characterized the conformational landscapes of the full-length S protein trimers detailing conformational transitions between functional states and unveiling previously unknown cryptic pockets [77]. A combination of computational protein design and cell-free expression functional pipeline to rapidly screen and optimize computationally designed mini-protein inhibitors of SARS-CoV-2 produced 75-residue ACE2 mimics that simultaneously engage all three RBDs and potently neutralize multiple VOC’s including Omicron (B.1.1.529) and Delta (B.1.617.2) with the greater potency than the monoclonal antibodies [78]. The emergence of the Omicron variant and immune escape can be determined by multiple fitness trade-offs balancing between tendency to evolve new mutations evading antibodies and maintain binding affinity for ACE2 [79–81].

Together, experimental and computational studies suggested that the cross-neutralization activity against Omicron variants and the nature of emerging escape mutation hotspots may be driven by balance and tradeoff of multiple factors such as protein stability, binding interactions, and the long-range allosteric interactions and communications in the S-RBD/antibody complexes. However, the dynamic and energetic details quantifying the balance and contribution of these factors, particularly the role of allosteric interactions and signaling into neutralization mechanisms of cross-reactive and ultra-potent antibodies against Omicron variant remain mechanistic and scarcely characterized. Our previous studies revealed that the SARS-CoV-2 S protein can function as an allosteric regulatory machinery that can exploit the intrinsic plasticity of functional regions controlled by stable allosteric hotspots to modulate specific regulatory and binding functions [82–87]. In this study, we combined all-atom MD simulations, the ensemble-based mutational scanning of protein stability and binding, and perturbation-based network profiling of allosteric interactions in the S-RBD and S-RBD Omicron structures with a panel of cross-reactive and ultra-potent single antibodies (B1-182.1 and A23-58.1) as well as combinations (A19-61.1/B1-182.1 and A19-46.1/B1-182.1). Using this approach, we quantify local and global effects of mutations in the S-RBD complexes, identify structural stability centers, characterize binding energy hotspots and predict the allosteric control points of long-range interactions and communications. A network-based perturbation approach for mutational profiling of allosteric residues potentials is proposed that evaluates the effect of antibody binding on modulation of allosteric interactions in the studied S-RBD complexes. This study shows that the local binding energetics of the antibodies and allosteric signaling in the complexes are tolerant to Omicron mutations. The results of this study suggest that immune evasion via mutations of the common hydrophobic stability hotspots may be restricted by the requirements of the RBD folding and binding to the ACE2 host receptor. We present evidence that mutation-sensitive and antibody-specific escape mutants may be selected to accommodate multiple fitness trade-offs between protein stability, conformational adaptability, binding specificity, and allosteric signaling that can determine the unique immune escape patterns without compromising spike activity and binding with the ACE2.

## 2. Materials and Methods

### 2.1. Molecular Dynamics Simulations

All structures were obtained from the Protein Data Bank [88]. During structure preparation stage, protein residues in the crystal structures were inspected for missing residues and protons. Hydrogen atoms and missing residues were initially added and assigned according to the WHATIF program web interface [89,90]. The missing loops in the studied cryo-EM structures of the SARS-CoV-2 S protein were reconstructed and optimized using template-based loop prediction approaches ModLoop [91], ArchPRED server [92] and further confirmed by FALC (Fragment Assembly and Loop Closure) program [93]. The side chain rotamers were refined and optimized by SCWRL4 tool [94]. The protein structures were then optimized using atomic-level energy minimization with a composite physics and knowledge-based force fields as implemented in the 3Drefine method [95]. The atomistic structures from simulation trajectories were further elaborated by adding N-acetyl glycosamine (NAG) glycan residues and optimized.

A series of 10 independent 500ns all-atom MD simulations were performed for each studied S-RBD complexes : S-RBD with A23-58.1 (pdb id 7LRS), S-RBD with B1-182.1 (pd id 7MLZ), S-RBD with A19-61.1/B1-182.1 (pdb id 7TBF), and S-RBD Omicron with A19-46.1/B1-182.1 (pdb id 7U0D) in an N, P, T ensemble in explicit solvent using NAMD 2.13 package [96] with CHARMM36 force field [97]. Long-range non-bonded van der Waals interactions were computed using an atom-based cutoff of 12 Å with switching van der Waals potential beginning at 10 Å. Long-range electrostatic interactions were calculated using the particle mesh Ewald method [98] with a real space cut-off of 1.0 nm and a fourth order (cubic) interpolation. SHAKE method was used to constrain all bonds associated with hydrogen atoms. Simulations were run using a leap-frog integrator with a 2 fs integration time step. Energy minimization after addition of solvent and ions was conducted using the steepest descent method for 100,000 steps. All atoms of the complex were first restrained at their crystal structure positions with a force constant of 10 Kcal mol^-1^ Å^-2^. Equilibration was done in steps by gradually increasing the system temperature in steps of 20K starting from 10K until 310 K and at each step 1ns equilibration was done keeping a restraint of 10 Kcal mol-1 Å-2 on the protein Cα atoms. After the restrains on the protein atoms were removed, the system was equilibrated for additional 10 ns. An NPT production simulation was run on the equilibrated structures for 500 ns keeping the temperature at 310 K and constant pressure (1 atm). In simulations, the Nose-Hoover thermostat [99] and isotropic Martyna-Tobias-Klein barostat [100] were used to maintain the temperature at 310 K and pressure at 1 atm respectively.

### 2.2. Distance Fluctuations Stability and Communication Analysis

We employed distance fluctuation analysis of the simulation trajectories to compute residue-based stability profiles. The fluctuations of the mean distance between each pseudoatom belonging to a given amino acid and the pseudo-atoms belonging to the remaining protein residues were computed. The fluctuations of the mean distance between a given residue and all other residues in the ensemble were converted into distance fluctuation stability indexes that measure the energy cost of the residue deformation during simulations [101–104]. The distance fluctuation stability index for each residue is calculated by averaging the distances between the residues over the simulation trajectory using the following expression:

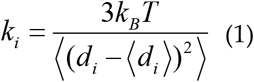

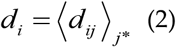

*d_ij_* is the instantaneous distance between residue *i* and residue *j, k_B_* is the Boltzmann constant, *T* =300K. 〈 〉 denotes an average taken over the MD simulation trajectory and *d_i_* = 〈*d_ij_*〉_*j*^*^_ is the average distance from residue *i* to all other atoms *J* in the protein (the sum over *J* implies the exclusion of the atoms that belong to the residue *i*). The interactions between the *C_α_* atom of residue *i* and the *C_α_* atom of the neighboring residues *i*−1 and *i*+1 are excluded in the calculation since the corresponding distances are constant. The inverse of these fluctuations yields an effective force constant *k_i_* that describes the ease of moving an atom with respect to the protein structure. The dynamically correlated residues whose effective distances fluctuate with low or moderate intensity are expected to communicate over long distances with the higher efficiency than the residues that experience large fluctuations. Our previous studies showed that residues with high value of these indexes often serve as structurally stable centers and regulators of allosteric signals, whereas small values of the distance fluctuation stability index are typically indicative of highly dynamic fluctuating sites [83,84]. The structurally stable and densely interconnected residues as well as moderately flexible residues that serve as a source or sink of allosteric signals could feature high value of these indexes.

### 2.3. Mutational Scanning and Sensitivity Analysis

We conducted mutational scanning analysis of the binding epitope residues for the SARS-CoV-2 S protein complexes. BeAtMuSiC approach [105–107] was employed that is based on statistical potentials describing the pairwise inter-residue distances, backbone torsion angles and solvent accessibilities, and considers the effect of the mutation on the strength of the interactions at the interface and on the overall stability of the complex. The binding free energy of protein-protein complex can be expressed as the difference in the folding free energy of the complex and folding free energies of the two protein binding partners:

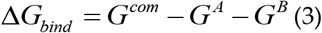

The change of the binding energy due to a mutation was calculated then as the following

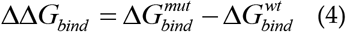

The binding free energy changes were computed by averaging the results over 1,000 equilibrium samples for each of the studied systems.

### 2.4. Network Analysis and Perturbation-Based Mutational Profiling of Allosteric Propensities

A graph-based representation of protein structures [108,109] is used to represent residues as network nodes and the inter-residue edges to describe non-covalent residue interactions. The network edges that define residue connectivity are based on non-covalent interactions between residue side-chains that define the interaction strength *I_j_* ac-cording to the following expression used in the original studies [108,109]:

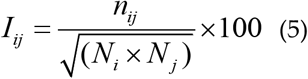

where *n_ij_* is number of distinct atom pairs between the side chains of amino acid residues *i* and *j* that lie within a distance of 4.5 Å. *N_i_* and *N_j_* are the normalization factors for residues *i* and *J*. We constructed the residue interaction networks using dynamic correlations [110] that yield robust network signatures of long-range couplings and communications. The ensemble of shortest paths is determined from matrix of communication distances by the Floyd-Warshall algorithm [111]. Network graph calculations were performed using the python package NetworkX [112]. Using the constructed protein structure networks, we computed the residue-based betweenness parameter. The short path betweenness centrality of residue *i* is defined to be the sum of the fraction of shortest paths between all pairs of residues that pass through residue *i*:

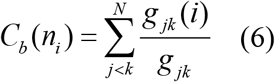

where *g_jk_* denotes the number of shortest geodesics paths connecting *J* and *k*, and *g_jk_*(*i*) is the number of shortest paths between residues *j* and *k* passing through the node *n_i_*. The betweenness centrality metric is also computed by evaluating the average shortest path length (ASPL) change by systematically removing individual nodes [113,114]. The following Z-score is then calculated

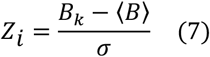

Through mutation-based perturbations of protein residues we compute dynamic couplings of residues and changes in the average short path length (ASPL) averaged over all modifications in a given position. The change of ASPL upon mutational changes of each node is inspired and reminiscent to the calculation proposed to evaluate residue centralities by systematically removing nodes from the network.

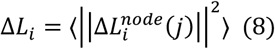

where *i* is a given site, *j* is a mutation and 〈⋯〉 denotes averaging over mutations. 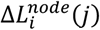 describes the change of ASPL upon mutation *j* in a residue node *i*. Δ*L_i_* is the average change of ASPL induced by all mutations of a given residue. Z-score is then calculated for each node as follows:

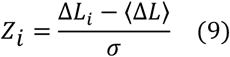

〈Δ*L*〉 is the change of the ASPL under mutational scanning averaged over all protein residues in the S-RBD and σ is the standard deviation. The ensemble-averaged Z–scores ASPL changes are computed from network analysis of the conformational ensembles using 1,000 snapshots of the simulation trajectory for the native protein system.

## 3. Results and Discussion

### 3.1. Structural Analysis of the S-RBD Complexes with Antibodies

We began with the structural analysis of the S-RBD binding with single antibodies A23-58.1 (Figure 1A-C) and B1-182.1 (Figure 1D-F). The cryo-EM structures of the S-RBD WA1 complexes [55] showed that both antibodies adapt the mode of binding by directly blocking the interaction of the RBD with ACE2 and could be classified as either class I (ACE2 blocking, binding RBD up only) or II (ACE2 blocking, binding RBD up or down) RBD antibodies [55]. A23-58.1 an B1-182.1 bind to an invariant region of the RBD tip with the binding epitopes residues K417, L455, F456, K458, Y473, Q474, A475,G476, N477, K478, C480, E484, G485, F486, N487, Y489 and Q493 (Figure 1A,D). Mapping of the binding epitope residues together with the sites of Omicron mutations (Figure 1B,E) highlighted a minor overlap where only K417, N477 and K478 positions engage in the interactions with the antibodies. The interaction details of the antibody-RBD interface showed that the RBD tip region binds to a “crater-like” region formed by the complementarity-determining regions (CDRs) of the antibodies (Figure 1C,F). The key intermolecular interactions are formed between the aromatic RBD residues F456, F486 and Y489 and a group of hydrophobic sites in the heavy and light chains of the antibodies. The most prominent contacts are mediated by F486 deeply protruding into the antibody “crater” and surrounded by P95, F100 (heavy chain) and Y91,G92, W96, Y32 (light chain) (Figure 1C,F). Other significant interactions are formed by Y489 of the RBD with the heavy chain sites C97, V52, A33 as well as by F456 of the RBD and the heavy chain positions T30, S31 S32,V52, G53, S54, (Figure 1C,F). Structural analysis of the antibody-RBD interfaces also highlighted contacts formed by Y473, A475 and N487 in the central interface area while other important sites T478 and Q493, which are among Omicron-targeted positions, are located on the periphery of the biding interface.

**Figure 1.**
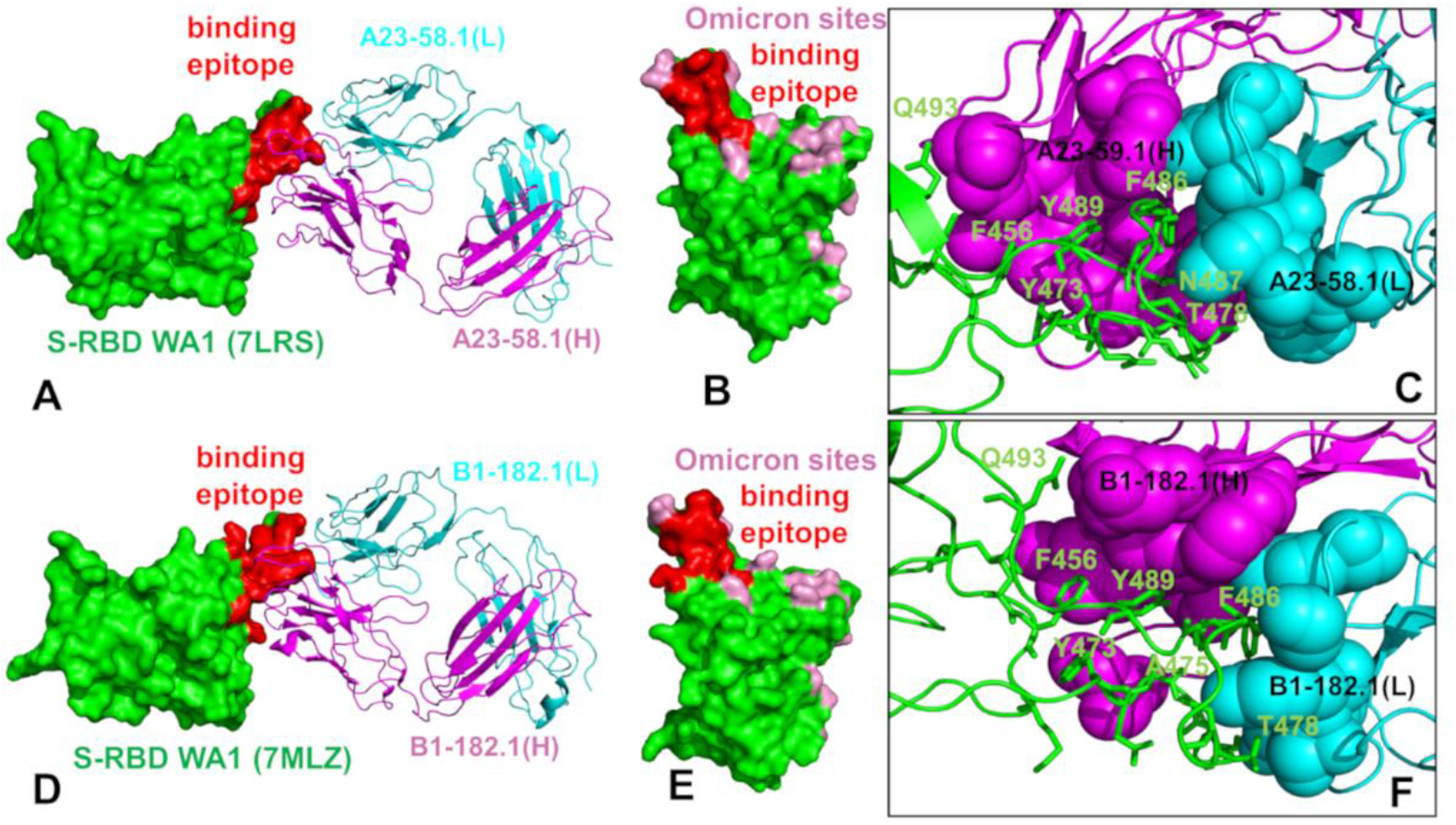
Structural organization of the SARS-CoV-2-RBD complexes with the A23-58.1 and B1-182.1 antibodies. (A) The cryo-EM structure of the S-RBD WA1 complex with A23-58.1. The S-RBD is in green surface, the binding epitope residues are colored in red, and the antibody is shown in ribbons. The heavy chain of A23-58.1 is in magenta and the light chain in cyan. (B) The S-RBD is shown in green surface with the binding epitope residues in red and the RBD sites of Omicron mutations (G339, S371, S373, S375, K417, N440, G446, S477, T478, E484, Q493, G496, Q498, N501, and Y505) are colored in pink. (C) A detailed closeup of the interacting residues in the binding interface of the S-RBD complex with A23-58.1. The S-RBD binding residues are shown in green sticks and annotated. The contact sites of the heavy antibody chain is in magenta spheres and in cyan spheres for the light chain residues. (D) The cryo-EM structure of the S-RBD WA1 complex with B1-182.1 antibody. The S-RBD is in green surface, the binding epitope residues are colored in red, and the antibody is shown in ribbons. The heavy chain of is in magenta and the light chain in cyan. (E) The S-RBD green surface with the binding epitope residues in red and the RBD sites of Omicron mutations in pink. (F) A closeup of the interacting residues in the binding interface of the S-RBD complex with B1-182.1. The S-RBD binding residues are shown in green sticks and annotated, and the contact sites on the antibody are shown in magenta and cyan spheres for the heavy and light chains respectively.

The structure of the S-RBD WA1 complex with A19-61.1/B1-182.1 antibodies [56] retained a similar binding mode for B1-182.1 and provided an extended binding epitope consisting of the RBD residues K417, L455, F456, K458, Y473, A475, G476, S477, T478, G485, F486, N487, C488, Y489, and Q493 (Figure 2A). The RBD residues involved in the contacts with A19-61.1 included a continuous stretch of residues 440-452 as well as F490, Q493, S494, G496 sites. A number of Omicron sites (N440, G446, K417, S477, T478 and Q493) belong to the extended binding epitope (Figure 2A,B). Structural mapping of the binding epitope illustrated the non-overlapping and complementary nature of the binding residues for A19-61.1 and B1-182.1 providing a significant coverage of the RBD regions and enabling an effective interference with the RBD-ACE2 binding [55,56]. A detailed inspection of the intermolecular interactions reinforced the conserved nature of the contacts between the RBD residues F456, F486 and Y489 and hydrophobic sites in the heavy and light chains of B1-182.1 (Figure 2C). The key RBD contacts with B1-182.1 are preserved in the S-RBD WA1 complex with A19-61.1/B1-182.1 and are similarly mediated by F486 (Figure 2C).

**Figure 2.**
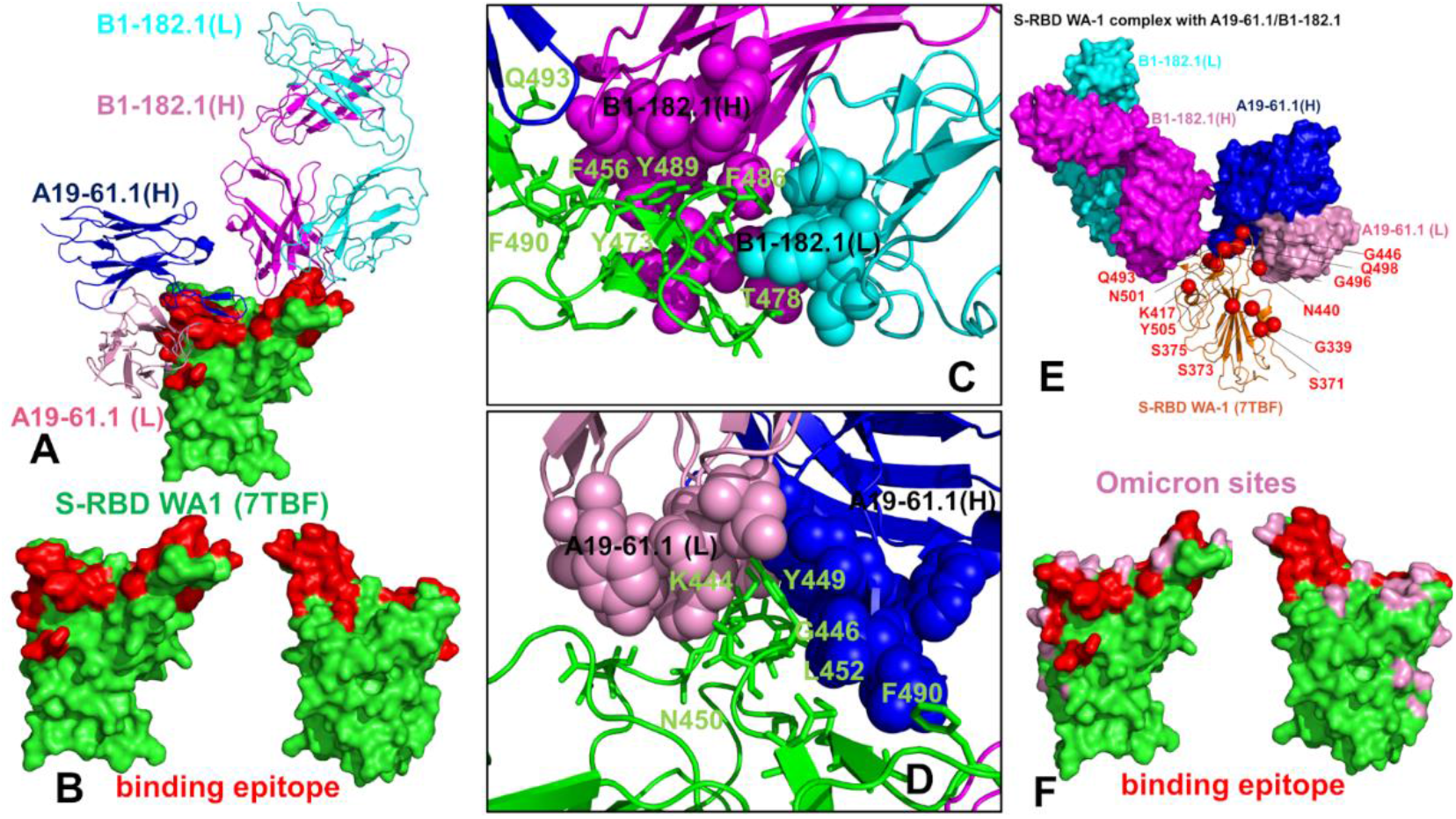
Structural organization of the SARS-CoV-2-RBD complex with the combination of A19-61.1/B1-182.1 antibodies. (A) The cryo-EM structure of the S-RBD WA1 complex with A19-61.1/B1-182.1 pair. The S-RBD is in green surface, the binding epitope residues are colored in red. The heavy chain of B1-182.1 is in magenta and the light chain in cyan. The heavy chain of A19-61.1 is in blue and the light chain is in pink. (B) The S-RBD is shown in two orientations in green surface with the binding epitope residues in red. (C,D) A closeup of the interacting residues in the binding interface of the S-RBD complex with B1-182.1 and A19-61.1 respectively. The S-RBD binding residues are shown in green sticks and annotated. The contact sites of the interacting antibodies are shown in spheres colored according to the established color scheme for the heavy and light chain. (E) The structure of the S-RBD WA1 complex with A19-61.1/B1-182.1 with the sites of Omicron mutations. The S-RBD is in orange ribbons, the Omicron sites are in red spheres and annotated. B1-182.1 is shown in surface (heavy chain in magenta, light chain in cyan) and A19-61.1 is in surface (heavy chain in blue, light chain in pink). (F) The S-RBD (green surface) with the binding epitope residues highlighted in red and sites of Omicron mutations in pink.

The peripheral interactions formed by T478 and Q493 residues are also maintained reflecting a considerable structural similarity of the S-RB WA1 complexes with a single B1-182.1 and A19-61.1/B1-182.1 combination. The binding interface of A19-61.1 is determined by K444, G446 and Y449 residues that penetrate into a cavity formed by the heavy chain residues H109, N112, I102, V104 and light chain residues S31, W32, D50 (Figure 2D). Noticeably, functionally important Q493 residue interacts with both B1-182.1 and A19-61.1 antibodies, suggesting a potential importance of this position in the stability of the complex and allostery. Indeed, Q493 forms interactions with G53, G55, N56 of the B1-182.1 heavy chain and A105, T107 of the A19-61.1 heavy chain (Figure 2C,D). Other important contacts with A19-61.1 are mediated by N450, L452, and F490 (Figure 2D). Structural maps of the binding epitope residues and sites of Omicron mutations in the complex (Figure 2E,F) showed that Omicron positions are generally either on the edges of the binding epitope or outside of the epitope regions with the exception of G446 position that is in the core of the binding interface.

The cryo-EM structure of the S-RBD Omicron with A19-46.1/B1-182.1 combination [56] revealed a consistently preserved binding mode of B1-182.1, while highlighting a more specific orientation and angle of approach for A19-46.1 antibody (Figure 3A,B). The binding epitope residues for B1-182.1 in the S-RBD Omicron complex with A19-46.1/B1-182.1 (Figure 3A,B) are essentially identical to those in the S-RBD WA1 complex with A19-61.1/B1-182.1 (Figure 2A,B). It can be noticed that the binding epitope for A19-46.1 has a substantial overlap with A19-61.1 but also featured a stretch of RBD residues 345-354, as well as residues D420, Y421, I468, S469, I470, E471, and A484 in the binding epitope that are specific for A19-46.1 (Figure 3A,B). The primary contacts formed by S-RBD Omicron with B1-182.1 are similarly mediated by hydrophobic sites F456, F486 and Y489, particularly between F486 and heavy chain positions A33, P95, D100 and light chain residues S31, Y32, Y91, W96 (Figure 3C). Additionally, the interaction contacts also engage Y473, N477, and K478 residues in binding with B1-182.1 (Figure 3C). The dominant contacts with the A19-46.1 antibody are mediated by the Y449 and F490 residues interacting with multiple sites in both heavy and light antibody chains (Figure 3D).

**Figure 3.**
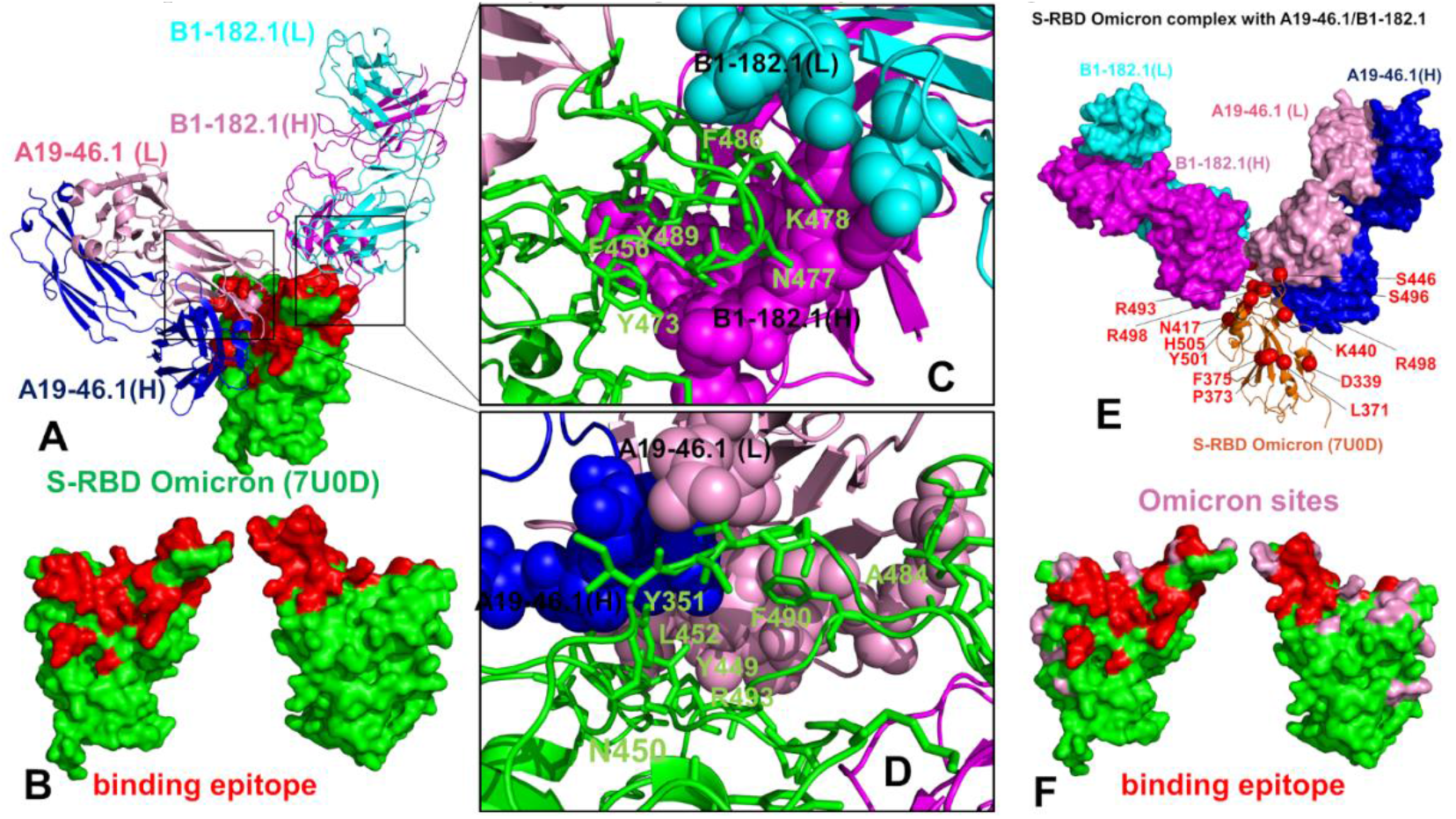
Structural organization of the SARS-CoV-2-RBD Omicron complex with the combination of A19-4.1/B1-182.1 antibodies. (A) The cryo-EM structure of the S-RBD WA1 complex with A19-46.1/B1-182.1 pair. The S-RBD is in green surface, the binding epitope residues are colored in red. The heavy chain of B1-182.1 is in magenta and the light chain in cyan. The heavy chain of A19-46.1 is in blue and the light chain is in pink. (B) The S-RBD is shown in two orientations in green surface with the binding epitope residues in red. (C,D) A closeup of the interacting residues in the binding interface of the S-RBD complex with B1-182.1 and A19-46.1 respectively. The S-RBD binding residues are shown in green sticks and annotated. The contact sites of the interacting antibodies are shown in spheres colored according to the established color scheme for the heavy and light chain. (E) The structure of the S-RBD WA1 complex with A19-46.1/B1-182.1 with the sites of Omicron mutations. The S-RBD is in orange ribbons, the Omicron sites are in red spheres and annotated. B1-182.1 is shown in surface (heavy chain in magenta, light chain in cyan) and A19-61.1 is in surface (heavy chain in blue, light chain in pink). (F) The S-RBD (green surface) with the binding epitope residues highlighted in red and sites of Omicron mutations in pink.

Additional interaction contacts feature RBD residues Y351, N450, L452, and Q493. The important new contacts formed by the S-RBD Omicron with A19-46.1 are established by the functionally important Omicron position A484 with the light chain residues Y33 and S26 of A19-26.2 (Figure 3D). Interestingly, a number of Omicron sites belong to or located on the edges of the extended binding epitope including K440, S446, K417, N477, K478, A484, R493, and S496 (Figure 3 E,F). Overall, structural analysis of the S-RBD WA1 and Omicron complexes with a panel of antibodies revealed a complex pattern of binding interactions and specific binding interfaces that are often dominated by non-polar interactions formed by a group of hydrophobic residues Y449, Y453, L455, A475, F456, F486, Y489 and F490 that form an important highly conserved patch that is important for the RBD stability and binding with the host receptor, and where mutations occur only at a very low frequency (< 0.05%) [63,64]. The structural analysis of these complexes provided a comprehensive map of the binding epitopes, but this is insufficient to assess which sites can be preferentially targeted by escape mutations and whether the pattern of escape mutants would be generic or antibody-specific.

The important question addressed in our study is how these cross-reactive ultra-potent antibodies manage to evade immune resistance from the mutated Omicron residues given a fairly noticeable presence of the Omicron positions in the binding epitope. In the next sections, we perform a comrehensive dynamic, energetic and network analysis of the antibody complexes to quantify at atomic level how the unique nature of the binding epitopes for these cross-reactive and ultrapotent antibodies allow for efficient binding while tolerating key residue substitutions in the Omicron sites and eliciting a mechanism of immune evasion with a limited and antibody-specific spectrum of mutant-escaping sites.

### 3.2. MD Simulations and Distance Fluctuation Analysis of Conformational Ensembles of the S-RBD Complexes with Antibodies Reveal Specific Dynamic Signatures and Stability Centers

All-atom MD simulations were performed for the S-RBD WA1 and S-RBD Omicron complexes with a panel of studied antibodies to examine the conformational landscapes and determine specific dynamic signatures induced by antibodies. Conformational dynamics profiles are described using the root mean square fluctuations (RMSF) obtained from simulations (Figure 4). The conformational mobility distribution for the S-RBD complex with A23-58.1 and B1-182.1 antibodies were similar, showing several deep local minima corresponding to residues 374-377, the RBD core residue cluster (residues 396-403), residues 445-456 that contain β-sheet β5 (residues 451-454) (Figure 4). We observed that the mobile flexible RBM loops (residues 473-487) become partly constrained in the complex owing to stabilizing contacts with B1-182.1 antibody that reduce local fluctuations in this region. Noticeably, residues 470-480 retain a certain degree of flexibility in the complex despite forming a portion of the intermolecular interface. At the same time, residues 481-495 and particularly key sites involved in the intermolecular contacts experience only minor fluctuations. The RBM residues 501-510 that are exposed to solvent showed more significant displacements. MD displayed a similar pattern of fluctuations in the S-RBD complex with A19-61.1/B1-182.1 (Figure 4). The most stable RBD positions corresponded to the RBD core residue clusters (residues 396-403 and 430-435). Interestingly, the RBD residues 440-452 involved in contacts with A19-61.1 showed minor fluctuations, while the binding epitope region 470-485 retained moderate mobility and the RBD binding interface sites 486-493 showed reduced fluctuations (Figure 4). The most interesting and intriguing pattern of the RBD dynamics was observed in the S-RBD Omicron complex with A19-46.1/B1-182.1. The dynamic profile showed a greater mobility of the S-RBD regions outside of the binding interface (residues 360-395). At the same time, the dynamic profile featured deep local minima corresponding to the specific regions, particularly residues 395-403, 420-423, 448-452, and 490-493 (Figure 4). By mapping positions of the Omicron mutations on the dynamics profiles, it could be seen that S371L, S373P and S375F positions experience more appreciable fluctuations, while the reduced mobility in K417N, N440 and G446S may be attributed to the stabilizing interactions with A19-46.1/B1-182.1 antibodies. Despite the intermolecular contacts, the RBM tip positions S477N, T478K remained moderately flexible but the Omicron positions Q493R and G496S involved in strong interactions with the antibodies correspond to well-defined local minima of the profile and become immobilized in the S-RBD Omicron complex (Figure 4). The important implication of these observations is that conformational plasticity of the S-RBD Omicron can be retained in the complex yielding a fairly narrow range of highly stabilized positions induced by binding which could potentially limit the emergence of resistant mutations.

**Figure 4.**
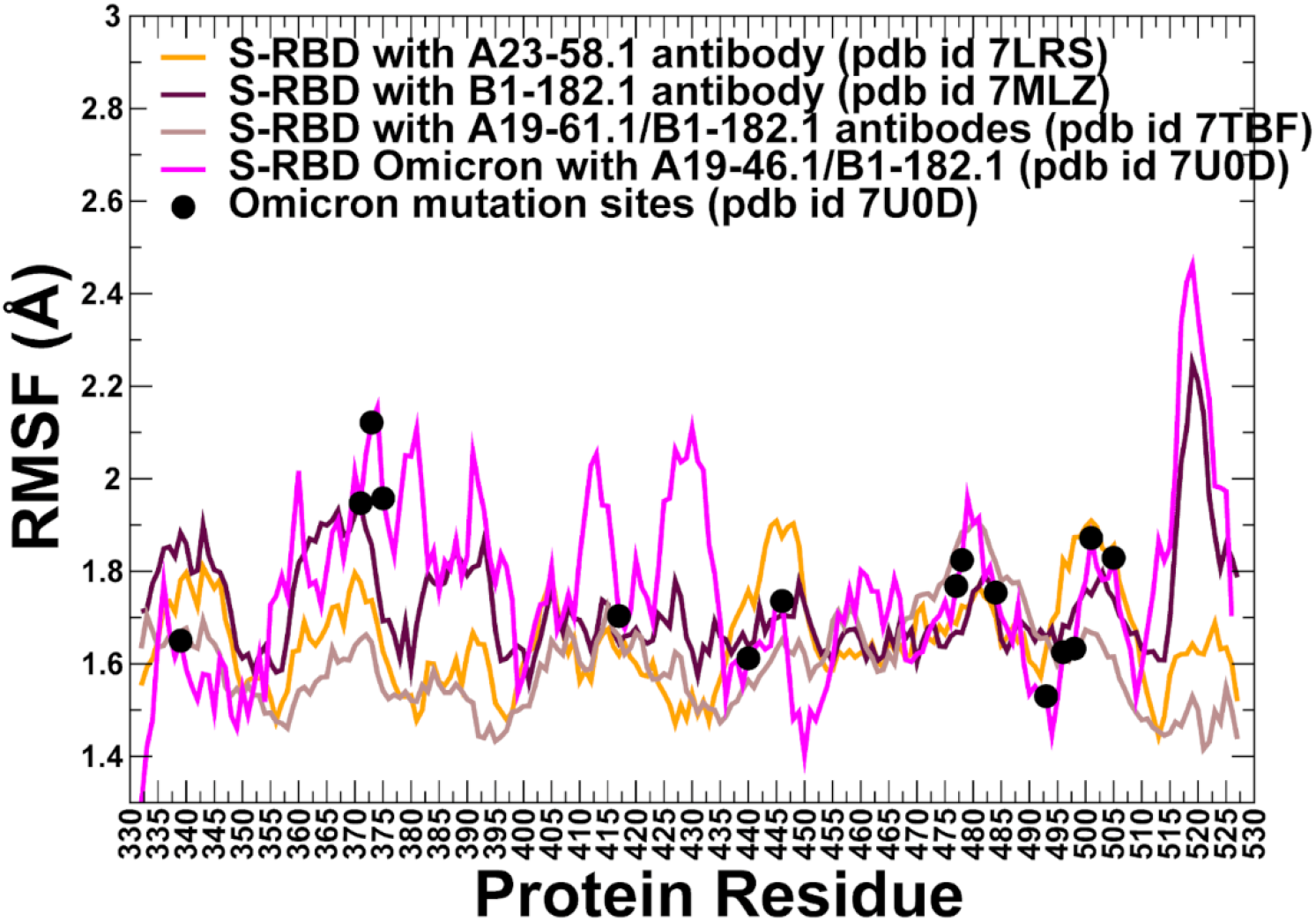
Conformational dynamics profiles obtained from simulations of the SARS-CoV-2 S-RBD complexes. The RMSF profiles for the RBD residues obtained from MD simulations of the S-RBD WA1 complex with A23-58.1 (orange lines), S-RBD with B1-182.1 (maroon-colored lines), S-RBD with A19-16.1/B1-182.1 (light brown lines), and S-RBD Omicron with A19-46.1/B1-182.1 (magenta colors). The positions of the Omicron sites are mapped on the dynamics profile of the S-RBD Omicron with A19-46.1/B1-182.1 and shown as black-filled circles.

Using conformational ensembles of the S-RBD complexes we computed the fluctuations of the mean distance between each residue and all other protein residues that were converted into distance fluctuation stability indexes that measure the energetics of the residue deformations (Figures 5,6). The high values of distance fluctuation indexes are typically associated with globally stable residues as they display small fluctuations in their distances to all other residues in the protein system, while small values of this parameter would correspond to the flexible sites that experience large deviations of their inter-residue distances. We first analyzed the distributions of the S-RBD WA1 complexes with A23-58.1 (Figure 5A,B) and B1-182.1 (Figure 5C,D). We observed several dominant and common peaks reflecting similarity of the topological and dynamical features of these S-RBD complexes. The RBD profiles for both complexes revealed a consistent pattern of local maxima that are aligned with the antibody-interacting positions K417, Y421, Y453, L455, F456, Y473, F486, N487 and Y489 (Figure 5A,C). Strikingly, the most dominant stability hotspots in the S-RBD complexes with A23-58.1 and B1-182.1 coreponded to the conserved hydrophobic positions L452, L455, F456, Y473, F486, Y489, F490 and L492 (Figure 5A,C). Our observations are in line with the structural analysis suggesting that A23-58.1, B1-182.1 and other broadly cross-reactive and ultrapotent antibodies can exhibit their activity by targeting this group of conserved and RBD stability-essential sites [59]. The stability peaks for the interacting antibodies A23-58.1 (Figure 5B) and B1-182.1 (Figure 5D) corresponded to the heavy chain positions T30, S31 S32 interacting with F456 on the RBD and the heavy chain residue cluster 90-95 that is anchored by P95 forming contacts with F486 position.

**Figure 5.**
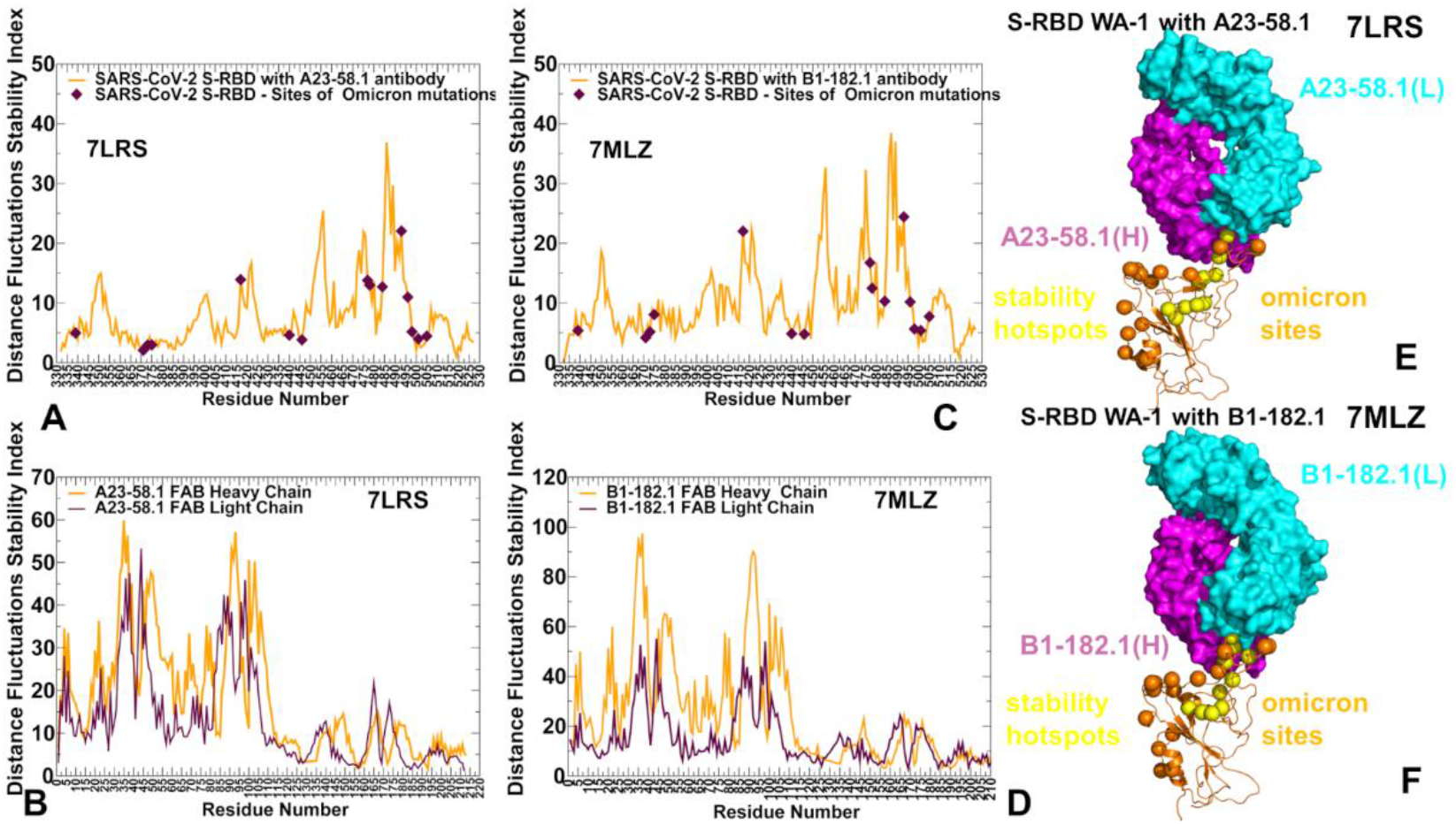
The distance fluctuation stability index profile obtained from simulations of the SARS-CoV-2 S-RBD complexes with A23-58.1 and B1-182.1 antibodies. (A) The stability index profile (shown in orange lines) for the RBD residues obtained from MD simulations of the S-RBD WA1 complex with A23-58.1 (pdb id 7LRS). The index values for the sites of Omicron mutations are highlighted by maroon-colored filled diamonds. (B) The distance fluctuation stability profile for the A23-58.1 antibody (heavy chain in orange lines and light chain in maroon lines). (C) The stability index profile (shown in orange lines) for the RBD residues obtained from MD simulations of the S-RBD WA1 complex with B1-182.1 (pdb id MLZ). The index values for the sites of Omicron mutations are in maroon-colored filled diamonds. (D) The stability profile for the B1-182.1 antibody (heavy chain in orange lines and light chain in maroon lines). (E,F) Structural maps of the stability centers for the S-RBD complexes with A23-58.1 and B1-182.1 antibodies respectively. The S-RBD is in orange ribbons. The heavy chains of the antibodies are shown in magenta surface and the light chains are in cyan-colored surface. The stability hotspots are shown in yellow-colored spheres. For comparison, the sites of Omicron mutations are highlighted in orange spheres.

**Figure 6.**
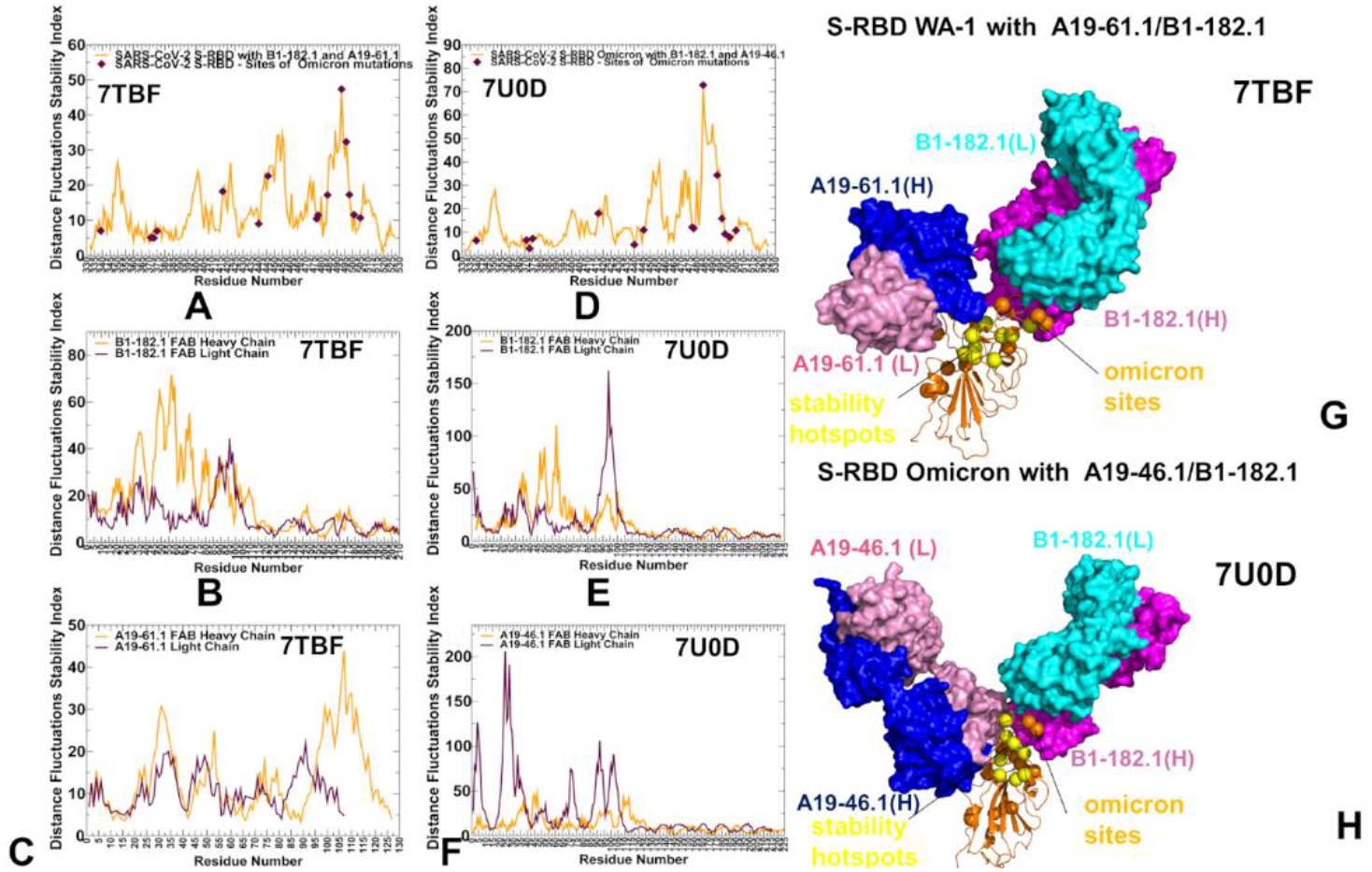
The distance fluctuation stability index profile obtained from simulations of the SARS-CoV-2 S-RBD complex with A19-61.1/B1-182.1 and S-RBD Omicron complex with A19-46.1/B1-182.1 (A) The stability index profile (shown in orange lines) for the RBD residues obtained from MD simulations of the S-RBD WA1 complex with A19-61.1/B1-182.1 (pdb id 7TBF). The index values for the sites of Omicron mutations are highlighted by maroon-colored filled diamonds. (B) The distance fluctuation stability profile for the B1-182.1 antibody in this complex (heavy chain in orange lines and light chain in maroon lines). (C) The distance fluctuation stability profile for the A19-61.1 antibody in this complex (heavy chain in orange lines and light chain in maroon lines). (D) The stability index profile (shown in orange lines) for the RBD residues obtained from MD simulations of the S-RBD Omicron complex with A19-46.1/B1-182.1 (pdb id 7U0D). The index values for the sites of Omicron mutations are in maroon-colored filled diamonds. (E) The stability profile for the B1-182.1 antibody in this complex (heavy chain in orange lines and light chain in maroon lines). (F) The stability profile for A19-46.1in this complex (heavy chain in orange lines and light chain in maroon lines). (G) Structural map of the stability centers for the S-RBD WA1 complex with A19-61.1/B1-182.1. (H) Structural map of the stability centers for the S-RBD Omicron complex with A19-46.1/B1-182.1. The S-RBD is in orange ribbons. The heavy chain of A19-61.1 and A19-46.1 is in blue surface and the light chain is in pink surface. The stability hotspots are shown in yellow-colored spheres and the sites of Omicron mutations are highlighted in orange spheres.

Structural map of the distribution peaks along with the Omicron positions (Figure 5E,F) illustrated the preferential localization of the stability centers in the RBD core and immobilized central region of the binding interface, while Omicron sites are distributed in the more flexible exposed regions of the RBD. Together, structural and dynamics analyses indicated that L455, F456, Y473, F486, N487 and Y489 residues become further rigidified in the complexes, largely due to their vital role in the binding interface (Figure 5A,B). These results are consistent with the functional experiments in which cell sur-face-expressed spike binding to A23-58.1 and B1-182.1 was knocked out by F486R, N487R, and Y489R mutations, completely abolishing neutralization [59]. Furthermore, for A23-58.1, several escape mutants include mutations of F486 particularly F486S mutation, while B1-182.1 escape can be mediated by F486L mutant [59]. Importantly, according to the original analysis [59], for A23-58.1 and B1-182.1, residues F486, N487, and Y489 were present in >99.96% of sequences, and only F486L was noted in the database at >0.01% (0.03%). The implications of the dynamics analysis revealing that these hydrophobic sites are associated with the pronounced stability hotspots suggests that the emergence of effective escape mutations in these sites may be limited as it would incur a significant stability fitness cost on the virus. Another important revelation of the distance fluctuation analysis is generally moderate or low stability indexes for the Omicron sites, reflecting conformational flexibility of these residues even for the Omicron positions ( S477, T478) involved in the interactions with the antibodies (Figure 5A,B). It is worth pointing out that only a single Omicron position Q493 displayed a moderate-high value of the stability index, indicating that substitutions in this site may potentially produce antibody-escaping mutants. Consistent with the predicted moderate stability effect, only marginal B1-182.1 escape can be mediated by Q493R but with minor impact on binding and no appreciable reduction of the neutralization activity [59]. According to this analysis the key stability hotpots of the S-RBD complexes corresponded to L452, L455, F456, Y473, F486, Y489, F490 and L492 positions, suggesting that mutations in these sites may weaken both local and long-range interactions and thus impair basic spike functionality. These findings are consistent with the recent stuies showing that residues Y449, L452, L455, E484, Y489, F490, L492, Q493, and S494 can be among immune-escaping hotspots that may destabilize binding with antibodies and erode neutralizing immune responses [115]. Furthermore, mutations of the common hydrophobic hotspots (Y449, Y473, L455, F456 and Y489) can disrupt both the stability of the RBD and binding to ACE2 and ultra-potent neutralizing antibodies [115]. These positions were identified in our analysis as critical stability hotspots of the S-RBD complexes. We argue that mutations of these hotspots can compromise the balance of multiple fitness trade-offs of the virus, specifically between immune escape the RBD stability and the affinity with the ACE receptor [79].

The distance fluctuation stability profile for the S-RBD complex with A19-61.1/B1-182.1 antibody combination showed an overall similar profile, reinforcing the notion that a subset of stability hotspots on the RBD may be preserved across S-RBD complexes (Figure 6A). The major stability profile peaks corresponded to the hydrophobic sites V350, V401, I402, L455, F456, Y473, A475, F486, and Y489 (Figure 6A). In addition, functional positions Q493 and S494 corresponded to the largest peak in the distance fluctuation profile, reflecting antibody-induced stabilization of these sites that are more flexible in the unbound form. Moreover, Q493 residue interacts with both B1-182.1 and A19-61.1 antibodies, and the increased structural stabilization of this site may be also associated with its potential mediating role in the long-range intermolecular interactions in the complex. Interestingly, several A19-61.1 interacting positions such as G446 and L452 were featured among moderate local peaks of the distribution (Figure 6A). In this context, it is worth noting that A19-61.1 escape mutations include G446V/S and S494R that occur at extremely low frequency [55,56] which is consistent with the role of these sites as structural stability hotspots in these complexes. The distance fluctuation profile for the B1-182.1 antibody showed strong peaks for residues 50-52 of the heavy chain (Figure 6B) that interacts with the RBD stability centers F486, Y489 and F490. The distribution for the A19-61.1 antibody displayed the largest peak for residues 107-112 of the heavy chain (Figure 6C) that interface with K444, G446, Y449, L452 positions on the RBD. The observed distribution patterns for the interacting antibodies A19-61.1 and B1-182.1 reflected stabilization of the key intermolecular interfaces and mediating role of the RBD stability centers G446, Y449, L452, F486, Y489, F490.

In the S-RBD Omicron complex with A19-46.1/B1-182.1 antibodies, there was some redistribution of the profile peaks detected in the N450, Y451, L452, L455, A484, and F490 positions (Figure 6D). Most of the distribution peaks are associated with the binding interface residues of A19-46.1 while the stability index values for the B1-182.1 interacting positions (Y473, L455, R493, N477, K478, G485, F486, N487, Y489) are more moderate (Figure 6D). Of interest is that Omicron mutational sites, with the exception of E484A and Q493R positions, displayed moderate stability indexes (Figure 6D). While E484A mutation can decrease stability of the unbound S-RBD, binding with A19-46.1 antibody in the S-RBD Omicron complex may favor this position as a mediating stability center (Figure 6D). This is in line with the functional data showing that E484A mutation did not affect neutralization capacity of A19-46.1 [56]. According to this structural investigation, E484A or Q493R substitutions result in complete loss of binding and neutralization activity for many classes I and II antibodies, whereas these mutations did not affect A19-46.1 neutralization and can be energetically favorable for antibody binding [56]. Strikingly, the experiments showed that escape mutations for A19-46.1 are generated in positions associated with the predicted stability hotspots N450, L452, and F490 (Figure 6D). Interestingly, among mutations mediating escape to A19-46.1 only F490L and L452R were present in the GISAID sequence database at greater than 0.01%. [55]. The distance fluctuation profile of the B1-182.1 antibody in the S-RBD Omicron complex displayed the highest peaks for the light chain residues L92, Y93, M94 that interface with A484 and F486 S-RBD sites (Figure 6E), while A19-46.1 antibody showed strong peaks for residues L24, S25, S26 of the light chain of the antibody that pack directly against A484 on the RBD (Figure 6F). In addition, some less significant stability peaks were seen for the heavy chain residues L105, L106, P107 of A19-46.1 (Figure 6F) packed against the RBD hotspots N450, Y451, and L452. Hence, in the S-RBD Omicron complex with A19-46.1/B1-182.1 one hotspot cluster is centered around A484 position interfacing with both antibodies, while several additional stability centers are mediated by N450, L452 and F490 positions that are engaged in binding with A19-46.1 antibody. It is worth noting that the distance fluctuation analysis showed the emergence of Q493/S494 and A484 positions as unique binding-induced stability centers respectively in the S-RBD complex and S-RBD Omicron complexes with synergistic antibody combinations (Figure 6). The relevance of these findings can be contextualized by comparison with the experimental studies showing that E484 and S494 are frequently engage in binding to antibodies but are more dispensable for binding to ACE2, and consequently are among hotspots of immune escape [117,118]. According to this study, mutations in S494 reduces neutralization competency of convalescent sera, and can facilitate broad antibody escape without significantly changing the binding affinity with the host receptor. Our data indicated that mutations in Q493 and S494 could potentially modulate stability and alter mediating capacity of these sites in the studied complexes. Indeed, Q493R mutation caused a 7-fold decrease of neutralization for B1-182.1 [55,56]. S494R mediates escape from A19-61.1 and A19-46.1 neutralization and is adjacent to other structurally stable sites Y449, N450 and L452 acting as hotspots mutational escape [55]. Another important finding of our analysis is that Omicron mutation sites generally featured small stability indexes, which indicates a substantial tolerance of structural stability across all studied complexes to Omicron modifications. This can contribute to the experimentally observed moderate impact of Omicron mutations on the efficiency of cross-reactive antibodies examined in our study. Structural mapping of stability centers and Omicron mutation sites (Figure 6G,H) highlighted only partial overlap between these groups. In particular, for the S-RBD Omicron complex with A19-46.1/B1-182.1 the structural projection of stability centers mimicked a “pathway” that connects the RBD core with the RBM residues serving as “bridges” between the synergistically acting antibodies. It may be argued that this structural disposition of stability centers is associated with their l role in mediating the long-range connectivity and communications in the complex. In some contrast, Omicron mutations are broadly distributed on the RBD interfaces and create a dynamic “shield” surrounding the stability regions that allows for modulation of the binding interactions with the antibodies.

Combined, the results of the conformational dynamics and distance fluctuation analysis showed that the binding-induced stability centers in the S-RBD complexes often corresponded to the antibody-specific immune-escaping hotspots. At the same time, the common stability hotspots shared among complexes include hydrophobic sites Y449, Y473, L455, F456 and Y489 that are constrained by the requirements for the RBD folding and binding with the ACE2 host receptor, and therefore may be limited in evolving antibody escaping mutants. These positions are also among the most targeted amino acids targeted by class I and II antibodies. Indeed, the most frequently targeted RBD residues by class I antibodies include L455, F456, F486, N487 and Y489, while for class II antibodies the most targeted sites are Y449, G485 and F486 [116]. Our findings suggested that the antibody-specific structural stability signatures of the S-RBD complexes can dictate the pattern of mutational escape and be one of the important contributing factors that determines immune escape hotspots.

### 3.3. Ensemble-Based Mutational Scanning and Energetic Cartography Identifies Binding Affinity Hotspots in the SARS-CoV-2 RBD Complexes with Ultrapotent Antibodies

By employing the conformational ensembles of the S-RBD WA1 and S-RBD complexes with antibodies we performed comprehensive mutational scanning of the interfacial RBD residues and computed binding free energy changes. In silico mutational scanning was done using BeAtMuSiC approach [105–107]. This approach allows for accurate predictions of the effect of mutations on both the strength of the binding interactions and the stability of the complex. The adapted in our study BeAtMuSiC approach was further enhanced through ensemble-based averaging of binding energy computations. The binding free energy ΔΔG changes were computed by averaging the results of computations over 1,000 samples obtained from simulation trajectories. To provide a comparison between the computational and experimental data, we constructed mutational heatmaps for the RBD binding interface residues (Figure 7). Intriguingly, despite structural similarities and common binding epitopes, binding heatmaps for the S-RBD WA1 complex with A23-58.1 (Figure 7A) and S-RBD Omicron complex with B1-182.1 (Figure 7B) displayed appreciable differences and featured several unique mutational signatures. In the S-RBD WA1 complex with A23-58.1 a fairly elevated level of mutational tolerance to binding interactions was observed for many epitope residues including K417, L455, Y473, A475, S477, T478, G485 and Q493 residues (Figure 7A). At the same time, the mutational heatmap clearly identified three major binding hotspots in F456, F486 and Y489 positions, where particularly large destabilization changes were induced by mutations of the F486 residue (Figure 7A). These results agree with the experimental data showing that binding and neutralization capacity of A23-58.1 can be markedly reduced by mutations in F486 and Y489 (particularly F486R/S, Y489R) and partly impaired by some mutations in F456 (F456R/Q/S) [55]. Consistent with the experiments, our energetic analysis showed that F486Q, F486R and F486S mutations yielded the most significant destabilization ΔΔG = 3.74 kcal/mol, ΔΔG = 3.75 kcal/mol and ΔΔG = 4.04 kcal/mol respectively (Figure 7A). According to the mutational scanning analysis, mutations in the Omicron sites S477, T478 and Q493 caused moderate destabilization changes with S477N, T478K and Q493R substitutions yielding ΔΔG = 0.14 kcal/mol, ΔΔG = 0.73 kcal/mol and ΔΔG =0.33 kcal/mol respectively for A23-58.1 antibody (Figure 7A). This is consistent with the observed tolerance of the antibody binding to mutations in these Omicron positions [55]. In general, the computed binding free energy changes reproduced the experimental data surprisingly accurately, confirming the validity of the energetic model [55,56].

**Figure 7.**
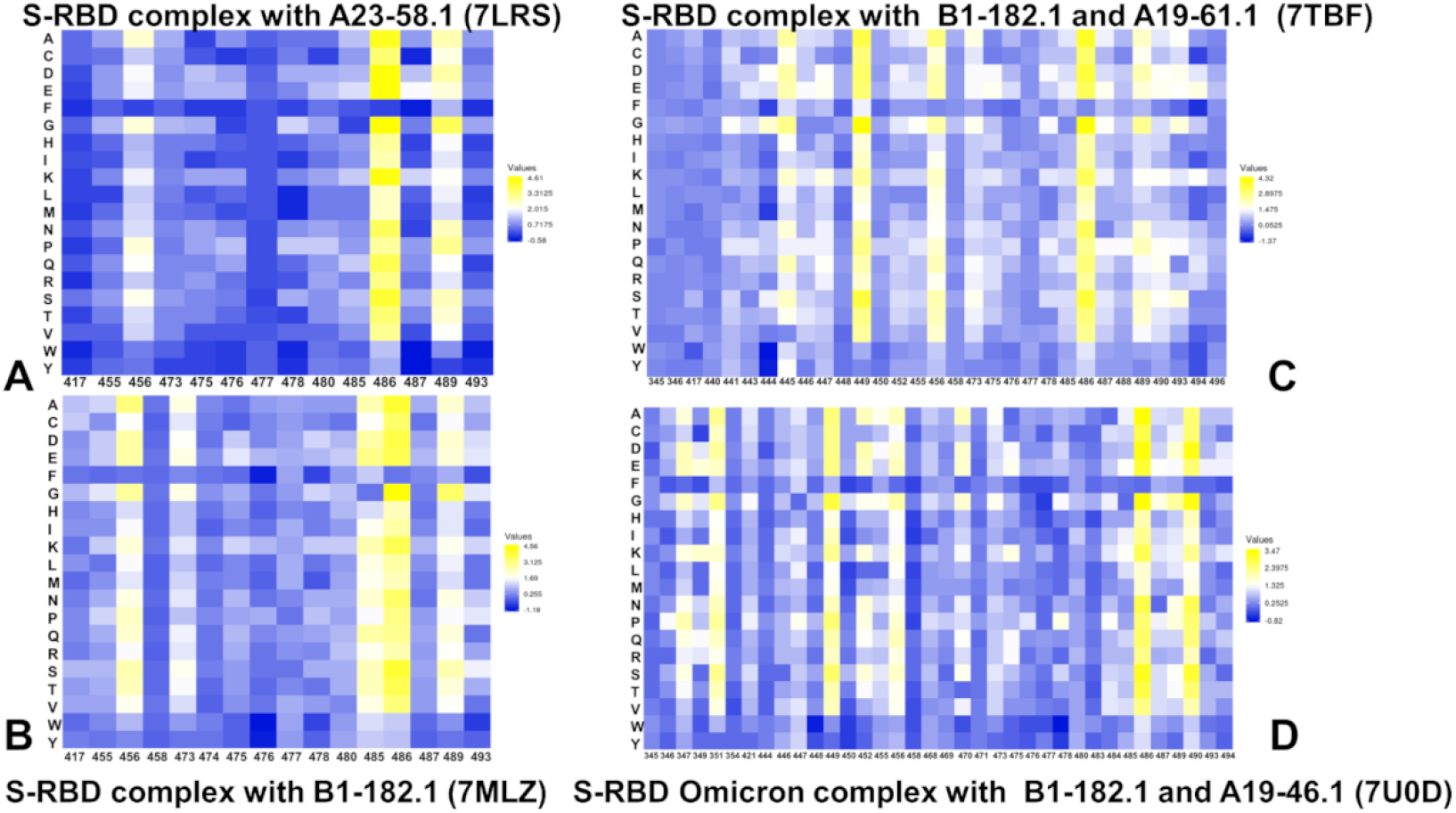
The ensemble-based mutational scanning of binding for the SARS-CoV-2 S-RBD and S-RBD Omicron complexes with ACE2. The mutational scanning heatmap for the binding epitope residues in the S-RBD WA1 complex with A23-58.1 (A), S-RBD WA1 complex with B1-182.1 (B), S-RBD WA1 complex with A19-61.1/B1-182.1 combination (C), and S-RBD Omicron complex with A19-46.1/B1-182.1 (D). The binding energy hotspots correspond to residues with high mutational sensitivity. The heatmaps show the computed binding free energy changes for 20 single mutations on the sites of variants. The squares on the heatmap are colored using a 3-colored scale blue-white-yellow, with yellow indicating the largest unfavorable effect on stability. The standard errors of the mean for binding free energy changes were based on a different number of selected samples from a given trajectory (500 and 1,000 samples) are within 0.07-0.16 kcal/mol.

The mutational heatmap of the S-RBD Omicron complex with B1-182.1 (Figure 7B) exhibited somewhat more sensitive profile of energetic changes, pointing to binding hotspots in F456, Y473, G485, F486, Y489, and Q493 positions. Common to both complexes, the most sensitive RBD positions corresponded to F456, F486 and Y489 sites (Figure 7A,B). However, we observed a greater binding sensitivity to mutations in F456 and Q493 sites. Our data revealed that F486Q, F486R and F486S mutations yielded the largest destabilization changes (ΔΔG = 2.95 kcal/mol, ΔΔG = 2.55 kcal/mol and ΔΔG = 3.92 kcal/mol respectively) that were moderately less detrimental as compared to their effect on binding with A23-58.1. Mutations of S477, T478 and Q493 similarly produced small energetic changes where Omicron mutations S477N, T478K and Q493R yielded ΔΔG = 0.26 kcal/mol, ΔΔG = 1.02 kcal/mol and ΔΔG = 0.60 kcal/mol respectively (Figure 7B). Structural maps of the binding epitope residues and hotspots for these S-RBD complexes (Figure 8A,B) illustrated a remarkable similarity in the binding modes and binding epitope residues. The structural mapping also highlighted differences between the binding energy hotspots and sites of Omicron mutations. It is evident that both A23-58.1 (Figure 8A) and B1.182.1 antibodies (Figure 8B) target a focused RBD tip region with only a small overlap with the Omicron positions.

**Figure 8.**
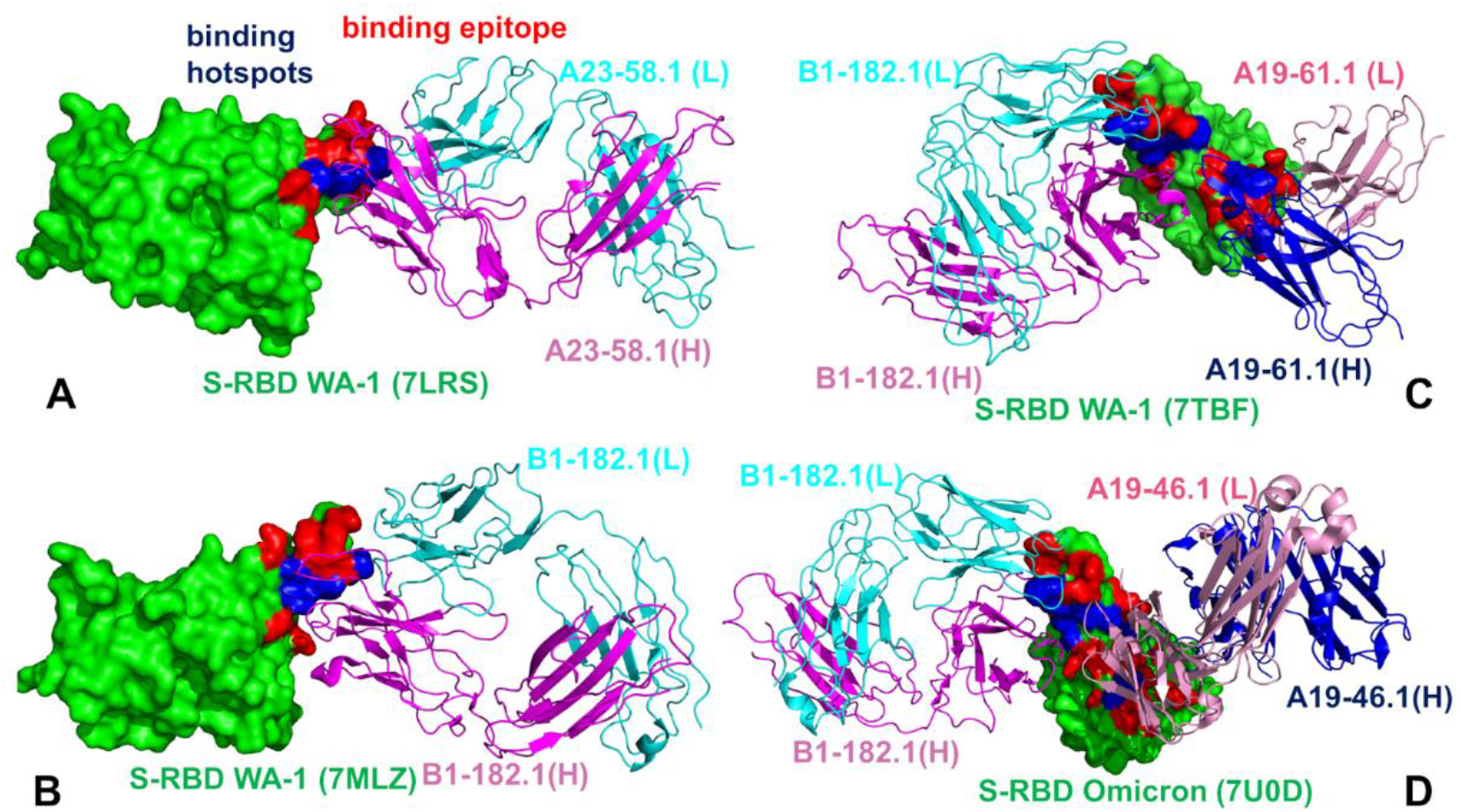
Structural maps of the binding epitope residues and binding energy hotspots in the S-RBD WA1 complex with A23-58.1 (A), S-RBD WA1 complex with B1-182.1 (B), S-RBD WA1 complex with A19-61.1/B1-182.1 combination (C), and S-RBD Omicron complex with A19-46.1/B1-182.1 (D). The S-RBD is in green surface, the binding epitope residues are colored in red, and the binding hotspots are colored in blue. The heavy chain of A23-58.1 and B1-182.1 on panels (A-D) is in magenta and the light chain in cyan. The heavy and light chains of A19-61.1 (panel C) and A19-46.1 (panel D) are in blue and pink ribbons respectively.

A more complex picture emerged from mutational heatmaps of the S-RBD complex with B1-182.1/A19-61.1 (Figure 7C) and S-RBD Omicron complex with B1-182.1/A19-46.1 (Figure 7D). In both complexes, the key binding hotspots corresponded to Y449, F456, F486, Y489 and F490 positions. Of particular significance is the fact that the conserved hydrophobic residues Y449, Y453, L455, F486, Y489 and F490 consistently emerged not only as the key stability centers but also as the dominant binding hotspots in the studied complexes (Figure 7). Structural maps of the binding interfaces for these complexes (Figure 8C,D) highlighted the extended binding epitope induced by the antibody pairs, also illustrating the binding hotspot clusters interacting with each of these antibodies. These findings are consistent with the recent stuies showing that these residues can destabilize binding with antibodies but at the unacceptable functional cost of disrupting the balance between various fitness tradeoffs, perturbing the intrinsic RBD stability and compromising binding to ACE2 [115]. This may narrow the “ evolutionary path” for the virus to adopt escape mutations in these key binding hotspots, thereby allowing for the cross-reactive antibodies to retain their neutralization efficiency. We also observed a strong binding sensitivity of A19-61.1 to mutations in G446 and Q493 positions, particularly Omicron mutations G446S, Q493R (Figure 7C). These results are in excellent agreement with the pioneering structure-functional experiments [55,56] that specifically emphasized the role of G446S mutant leading to a complete loss in activity for A19-61.1 against Omicron variant. The energetic heatmap analysis of the S-RBD Omicron complex with A19-46.1/1-182.1 detected an appreciable mutational sensitivity for residues K444, S446, N450, L452, and F490 (Figure 7D). In addition, we found that mutations in the A484 position which is an important mediating center are moderately destabilizing for the binding interactions. It is worth underscoring that the computed destabilization binding changes for the known escape mutations Y449S, N450S, N450Y, and L452R [55] were less dramatic as compared to mutations of the major binding hotspots.

These findings suggested that a mechanism of immune evasion and the escape mutants for cross-reactive antibodies may not be solely driven by the binding interactions but rather be determined by a combination and balance of different contributing local and global factors such as mutational effects on structural stability, binding strength, allosteric signaling and long-range communications. In general, class II antibodies can be often escaped by mutations at sites L452, E484, F490, and Q493 [115] suggesting that immune evasion mechanisms may operate by employing and weighting a multitude of contributing factors that affect virus fitness.

### 3.4. Allosteric Mutational Profiling of the Interaction Networks in the S-RBD Complexes Discern Sites and Mechanisms of Mutational Escape

We performed dynamic network analysis of the conformational ensembles and employed a recently introduced perturbation-based network approach for mutational scanning of allosteric residue potentials [87] to characterize allosteric interaction networks and identify allosteric hotspots in the S-RBD complexes. In the proposed model, allosteric hotspots are identified as residues in which mutations incur significant edgetic perturbations of the global residue interaction network that disrupt the network connectivity and cause a significant impairment of global network communications. Using a graph-based network model of residue interactions [108,109] in which network edges between nodes are weighted using dynamic residue-residue correlations obtained from MD simulations [110], we computed the ensemble-averaged distributions of several residue-based topological network metrics (Figures 9,10). The short path residue centrality (SPC) is used to analyze modularity and community organization of the dynamic residue interaction networks. The SPC distributions reflect the extent of residue connectivity in the interaction networks and allow for characterization of the mediating clusters in the complexes. The Z-score betweenness centrality is based on computing the average shortest path length (ASPL) as outlined in detail in the Methods section. By systematically introducing mutational changes in the S-RBD positions and using the equilibrium ensemble of the original system, we reevaluate the dynamic inter-residue couplings and compute mutation-induced changes in the ASPL parameter. These changes are then averaged over all substitutions in a given residue. In this manner, we characterize the average mutational sensitivity of each residue node on the changes in the network modularity and allosteric communications. By identifying residues where mutations on average induce a significant increase in the ASPL metric and therefore have a dramatic effect on the allosteric interaction network, we locate allosteric control points and regulatory hotspots that control long-range communications in the S-RBD complexes.

**Figure 9.**
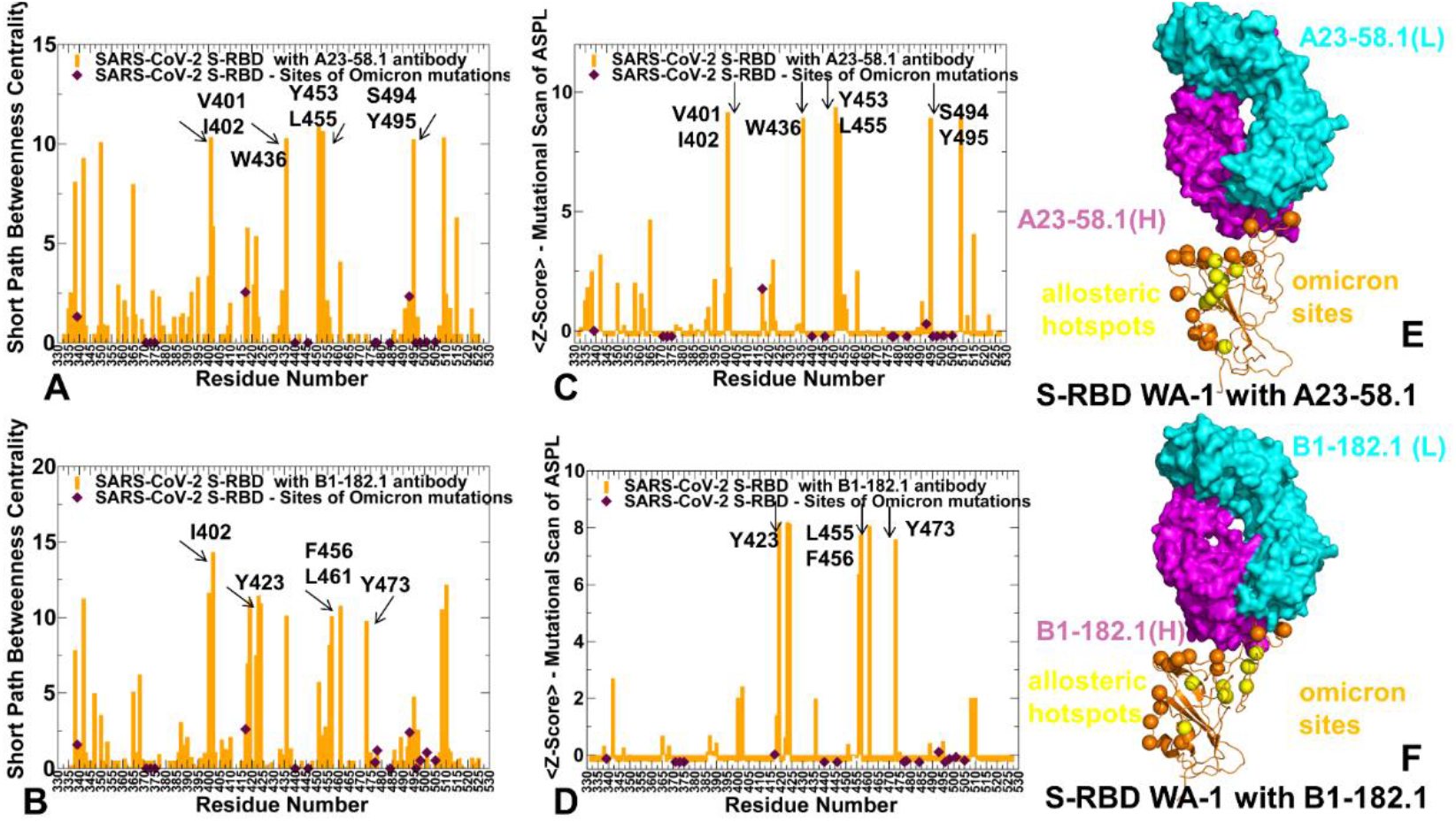
The network-based analysis of the S-RBD complexes with A23-58.1 and B1-182.1 anti-bodies. (A) The SPC centrality profile (shown in orange bars) for the RBD residues in the S-RBD WA1 complex with A23-58.1 (pdb id 7LRS). (B) The SPC centrality profile (shown in orange bars) for the RBD residues in the S-RBD WA1 complex with B1-182.1 (pdb id 7MLZ). The SPC values for the sites of Omicron mutations are highlighted by maroon-colored filled diamonds. (C) The Z-score ASPL profile (shown in orange bars) for the RBD residues in the S-RBD WA1 complex with A23-58.1. (D) The Z-score ASPL profile for the RBD residues in the S-RBD WA1 complex with B1-182.1. (E,F) Structural mapping of the allosteric centers associated with the Z-score ASPL peaks for the S-RBD complexes with A23-58.1 and B1-182.1 respectively. S-RBD is in orange ribbons. The heavy chain for the antibodies is in magenta surface and the light chain is in cyan-colored surface. The allosteric hotspots are in yellow spheres and sites of Omicron mutations are in orange spheres.

**Figure 10.**
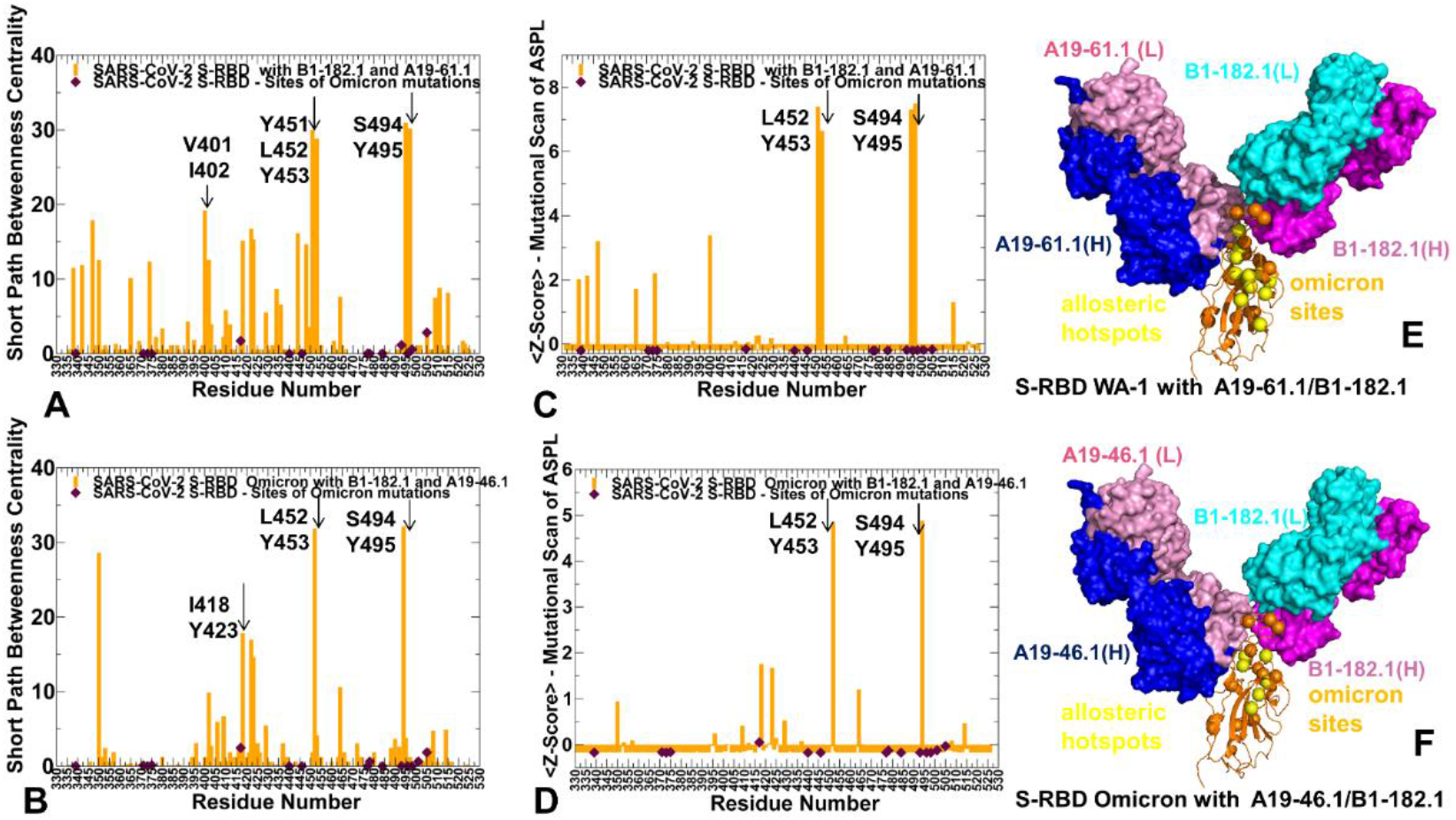
The network-based analysis of the S-RBD WA1 and S-RBD Omicron complexes with antibody combinations. (A) The SPC centrality profile (shown in orange bars) for the RBD residues in the S-RBD WA1 complex with A19-61.1/B1-182.1 (pdb id 7TBF). (B) The SPC centrality profile (shown in orange bars) for the RBD residues in the S-RBD Omicron complex with A19-46.1/B1-182.1(pdb id 7MLZ). The SPC values for the sites of Omicron mutations are highlighted by maroon-colored filled diamonds. (C) The Z-score ASPL profile (shown in orange bars) for the RBD residues in the S-RBD WA1 complex with A19-61.1/B1-182.1. (D) The Z-score ASPL profile for the RBD residues in the S-RBD Omicron complex with A19-46.1/B1-182.1. (E,F) Structural mapping of the allosteric centers associated with the Z-score ASPL peaks for the respective S-RBD complexes. The allosteric hotspots are shown in yellow spheres and sites of Omicron mutations are in orange spheres. S-RBD is shown in orange ribbons. The heavy chain for B1-182.1 is in magenta surface and the light chain is in cyan-colored surface. The heavy chain for A19-61.1 and A19-46.1 is in blue surface and light chain is in pink surface on panels E and F respectively.

The SPC distribution for the S-RBD complexes with A23-58.1 (Figure 9A) and B1-182.1 antibodies (Figure 9B) revealed dense clusters of high centralities in the regions that are often aligned with the stability centers. The emergence of local clusters of mediating residues implies a high level of connectivity in the residue interaction network which may allow for diverse communication routes in the complex. The high centrality peaks in both S-RBD complex with A23-58.1 corresponded to the hydrophobic sites V350, V401, I402, W436, Y453, L455, S494, and Y495 (Figure 9A). A similar but less dense distribution is seen in the S-RBD complex with B1-182.1 (Figure 9B). The key peaks in this distribution are I402, Y423, F456, L461, and Y473 residues. Interestingly, the mediating clusters of long-range interactions in these complexes are anchored by a group of residues that only partly overlap with the stability and binding hotspots. We noticed that Omicron positions K417 and Q493 displayed moderate SPC values, indicating that these sites can be also involved in mediating allosteric interactions. Perturbation-based mutational scanning of allosteric residue propensities provided information about potential allosteric hotspots by mapping a space of network-altering allosteric ‘edgetić variant sites (Figure 9C,D). In this model, the peaks of the Z-score ASPL profile corresponded to sites where mutational changes can incur a significantly increased ASPL values which implies a dramatic loss in the network connectivity. Importantly, the major peaks of the SPC and Z-ASPL distributions correspond to the same group of key mediating sites, but the ASPL profile allows to clearly distinguish mutation-sensitive hotspots of allosteric interactions.

Indeed, in the S-RBD complex with A23-58.1 the key allosteric hotspots revealed by the Z-score ASPL profile were aligned with I402, W436, Y453, L455, S494, and Y495 positions (Figure 9C) while in the complex with B1-182.1 antibody, the major peaks singled out Y423, L455, F456, L461 and Y473 residues (Figure 9D). In network terms, mutations in these positions could affect multiple intra and inter-molecular interactions altering the network connectivity which could adversely affect the long-range allosteric interactions. This analysis revealed several antibody-specific allosteric centers such as Y453, S494 and Y495 in the complex with A23-58.1 while L461 and Y473 in the complex with B1-182.1 antibody. Notably, the sites of Omicron mutations in these S-RBD complexes featured small Z-score ASPL values in both complexes, indicating that allosteric communications between S-RBD and antibodies may be tolerant to modifications in the Omicron positions (Figure 9C,D). Structural projection of the allosteric centers showed that the RBD positions mediating network connectivity could form a “pathway” connecting the RBD core with the central part of the binding interface (Figure 9E,F). The Omicron sites occupy more flexible regions and are dynamically coupled with the stable allosteric centers.

In the S-RBD complex with A19-61.1/B1-182.1, the shape of the SPC centrality profile is similar, but the most allosterically sensitive to mutations positions corresponded to Y451, Y453, S494, and Y495 residues (Figure 10A). These peaks were further accentuated in the Z-score ASPL profile (Figure 10B), indicating that mutations of these residues may perturb network connectivity and alter allosteric interactions in the S-RBD complex. Notably, among these allosteric hotspots only belongs to the binding epitope but the binding free energy changes induced by S494 mutations are moderate and comparable to binding effects of the other epitope sites (Figure 7C). At the same time, our results showed that S494 is among structural stability centers and is a specific allosteric hotspot in the complex with A19-46.1/B1-182.1. The experimental data showed that mutations in S494 (particularly S494R and S494P) can mediate escape from A19-61.1 [55], which according to our analysis may result not only from the moderate binding losses but mainly due to deleterious effect on the allosteric signaling in the complex. S494 and Y495 may be vulnerable target for escape mutations across multiple antibodies due their moderate effect on ACE2 binding [119].

For the S-RBD Omicron complex with A19-46.1/B1-182.1 we found that N450, L452 and S494 are the dominant peaks of the SPC and ASPL distributions (Figure 10C,D) and correspond to potential allosteric hotspots of the interaction network in which mutations can markedly alter efficiency of long-range interactions. These positions are also binding epitope residues. While mutational scanning showed that substitutions of these sites induce appreciable binding affinity losses, the binding free energy changes for the experimentally observed escape mutations N450S, N450Y, and L452R [55] are not markedly different from mutation-induce destabilization in other epitope sites. We argue that the observed functional effect of escape mutants may be determined by the cumulative contribution of these substitutions on the binding interactions, stability and allosteric signaling. Similarly, the sites of Omicron mutations exhibited small network centrality and ASPL values. Hence, the efficiency of binding and allosteric communications may be tolerant to the Omicron mutations. Structural mapping the allosteric centers highlighted the proximity of the stable allosteric centers and Omicron sites, also illustrating the connectivity of the allosteric hotspots linking the RBD core with the intermolecular interfaces (Figure 10E,F).

## 4. Conclusions

In this study, we combined all-atom MD simulations, the ensemble-based mutational scanning of protein stability and binding, and perturbation-based network profiling of allosteric interactions in the S-RBD and S-RBD Omicron structures with a panel of cross-reactive and ultra-potent single antibodies (B1-182.1 and A23-58.1) as well as combinations (A19-61.1/B1-182.1 and A19-46.1/B1-182.1). Using this approach, we quantify local and global effects of mutations in the S-RBD complexes, identify structural stability centers, characterize binding energy hotspots and predict the allosteric control points of long-range interactions and communications. By employing an integrated analysis of conformational dynamics and binding energetics, we establish that the examined ultra-potent antibodies against Omicron variant primarily target the conserved stability centers Y449, Y453, L455, F486, Y489 and F490 that also the dominant binding hotspots in the studied complexes. Th results suggested that mutations of these residues can severely impair binding with antibodies but at the unacceptable functional cost of disrupting the intrinsic RBD stability and compromising binding to ACE2. This may narrow the “ evolutionary path” for the virus to adopt escape mutations in these key binding hotspots, thereby allowing for the cross-reactive antibodies to retain their neutralization efficiency. The results revealed that protein stability and binding of the cross-variant potent antibodies are insensitive to the Omicron mutations, which is consistent with structural and biophysical experiments. Perturbation-based mutational scanning of allosteric residue propensities provided information about potential allosteric hotspots by mapping a space of network-altering allosteric ‘edgetic’ variant sites. By identifying residues where mutations on average induce a significant increase in the ASPL metric and therefore have a dramatic effect on the allosteric interaction network, we locate allosteric control points and regulatory hotspots that control long-range communications in the S-RBD complexes. Strikingly, these sites are different from the structural stability centers and binding hotspots. We found that some antibody-specific allosteric centers mediating long-range interactions correspond to important sites of mutational escape N450, L452 and S494 while only moderately affecting the RBD-antibody binding. These findings suggested that a mechanism of immune evasion that is determined by a combination and balance of different contributing local and global factors such as mutational effects on structural stability, binding strength, allosteric signaling and long-range communications. The insights from this investigation suggest therapeutic venues for targeted exploitation of the binding hotspots and allosteric communication centers that may potentially aid in evading drug resistance.

## Author Contributions

Conceptualization, G.V.; methodology, G.V.; software, G.V.; validation, G.V.; formal analysis, G.V.; investigation, G.V.; resources, G.V.; S.A.; K.K.; R.K.; data curation, G.V.; writing–original draft preparation, G.V.; writing–review and editing, G.V.; S.A.; K.K.; R.K.; visualization, G.V.; supervision, G.V.; project administration, G.V.; funding acquisition, G.V. All authors have read and agreed to the published version of the manuscript.

## Institutional Review Board Statement

Not applicable.

## Informed Consent Statement

Not applicable.

## Data Availability Statement

The data presented in this study are fully described in the manuscript

## Acknowledgments

The author acknowledges support from Schmid College of Science and Technology at Chapman University for providing computing resources at the Keck Center for Science and Engineering.

